# De novo design of α-helical peptide channels with designer stoichiometry

**DOI:** 10.1101/2024.10.05.616771

**Authors:** Ai Niitsu, Andrew R Thomson, Alistair J Scott, Jason T Sengel, Jaewoon Jung, Kozhinjampara R Mahendran, Mikiko Sodeoka, Hagan Bayley, Yuji Sugita, Derek N Woolfson, Mark I Wallace

## Abstract

Despite advances in peptide and protein design, the rational design of membrane-spanning peptides that form conducting channels remains challenging due to our imperfect understanding of the sequence-to-structure relationships that drive membrane insertion, assembly, and conductance. Here, we describe the design and computational and experimental characterization of a series of coiled coil-based peptides that form transmembrane α-helical barrels.

Through a combination of rational and computational design, we obtain barrels with 5 to 7 helices, as characterized in detergent micelles. In lipid bilayers, these peptide assemblies exhibit two conductance states with relative populations dependent on the applied potential: (i) a low-conductance states that correlate with variations in the modeled coiled-coil barrel geometries, indicating stable transmembrane α-helical barrels; and (ii) high-conductance states in which single pores change size in discrete steps. Notably, the high-conductance states are similar for all peptides in contrast to the low-conductance states. This indicates the formation of large, dynamic pores through the recruitment and expulsion of peptides, as observed in natural barrel-stave peptide pores. These findings establish rational routes to design and tune functional membrane-spanning peptide channels with specific conductance and geometry.

## Introduction

The *de novo* design of membrane-spanning peptides and proteins is challenging. While there have been recent successes in minimal,^1^ rational,^2^ and computational peptide and protein designs,^3–6^ the number of such designs is dwarfed by growing successes in the *de novo* design of water-soluble peptide assemblies and protein structures.^7^ This distinction arises from the increased technical difficulties of modeling, isolating, working with, and characterizing hydrophobic, membrane-soluble peptides and proteins. The primary problem in the *de novo* design of membrane-spanning biomolecules is the deficiencies in our understanding of how structure translates into function responses.^7–9^ In short, we need better design methods to define design targets, better overall design rules, and better ways to specify them over alternative states. Designing the stoichiometry of subunits in peptide-based assemblies would be a key step towards constructing more-complex structures and functions.^10^

α-Helical barrels (αHBs) are a promising architecture for the *de novo* design of functional membrane-spanning structures. αHBs are a class of coiled-coil assembly, which are well-understood protein structures that have been widely exploited in the design of water-soluble peptides and proteins.^10, 11^ In these structures, a defined number (‘n’) of α helices are arranged around a central channel with C_n_ or D_n/2_ symmetry, *i.e.*, parallel or antiparallel arrangements of adjacent helices, respectively. Distinctively, neighboring helices interact intimately via knobs-into-holes (KIH) packing of side chains, which is the structural signature of coiled coils. Although most natural coiled coils are found in water-soluble or fibrous proteins, for αHBs there are membrane-spanning and membrane-associated proteins for which structures have been elucidated; *e.g.*, pentamers (phospholamban,^12^ CorA),^13^ a heptamer (putative Plasmodium translocon of exported proteins),^14^ an octamer (Wza),^15^ and a dodecamer (Na^+^-driven membrane rotor ring of a bacterial ATP synthase).^16^ Inspired by these natural examples, peptide-based membrane-spanning αHBs have also been engineered,^17–19^ illustrating the utility of a strategies based on optimizing helix–helix packing.

For example, the repacking of phospholamban renders stable and fully defined pentamers.^20^ Computational design has been used to generate protein pores TMHC6 and TMH4C4 with 12 and 16 helices, respectively, organized in two concentric barrels.^3^ And previously, we have designed and produced self-assembling coiled-coil peptide ion channels that readily insert into lipid membranes.^2^ These designs are either derived directly from, or employ, sequence-to-structure relationships emanating from computationally designed, water-soluble αHBs.^11^ These strategies demonstrate the importance of optimizing helix–helix packing in both water-and membrane-soluble peptide assemblies and proteins to maintain stable αHBs. These successes also highlight the challenges in controlling the stoichiometry and stability of helical assemblies in membranes. These difficulties arise from the competition for hydrophobic residues to participate in both helix-helix and helix-membrane interfaces. Recently, the accessibility and accuracy of protein-structure prediction have been improved significantly for both water-soluble and membrane proteins through deep-learning methods such as AlphaFold.^21–24^ These methods are also being applied successfully to protein assemblies and oligomers.^25–27^ Nonetheless, predicting homo-oligomeric peptide structures without prior knowledge of their stoichiometry is still challenging. This is because of the lack of information linking amino-acid sequence patterns and specific oligomer states, and the small energetic differences between similar oligomers, exacerbated by the aforementioned competition for hydrophobic residues. For instance, amphipathic membrane-peptide assemblies can be non-specific and membrane-disruptive, as is typical of most antimicrobial peptides.^28^

To address these challenges, this study focuses on sequence-to-oligomer state relationships in membrane-spanning peptide αHBs. Using a combination of rational sequence design and computational *de novo* design, we optimize the steric packing of coiled-coil helices with the aim of generating pentameric, hexameric, and heptameric αHBs to insert spontaneously into membranes and form ion-conductive channels. We characterize these designs both experimentally and computationally leading to a relationship between the designed coiled-coil geometries and channel conductance.

## Results

### Computational design of coiled-coil packing yields new transmembrane αHB sequences

Most coiled coils are based on 7-residue (heptad) sequence repeats, which define the KIH interactions that specify and stabilize coiled-coil assemblies. The positions with the heptad repeat are conventionally labeled ***a – g***, with the ***a*** and ***d*** sites forming a seam that defines a single helix-helix interface as found in canonical dimeric coiled coils (Figure 1a). However, for αHBs (which have 5 or more helices) this is more complicated as the helix-helix interfaces and the resulting seams of interacting residues are wider. As a result, for these so-called Type-II coiled-coil interfaces,^3^ effectively two helical seams abut with an interface angle ∼103°, and side chains at the ***g***, ***a***, ***d***, and ***e*** positions all engage in the helix-helix interactions. Thus, all of these positions must be designed to specify αHBs.

**Figure 1.**
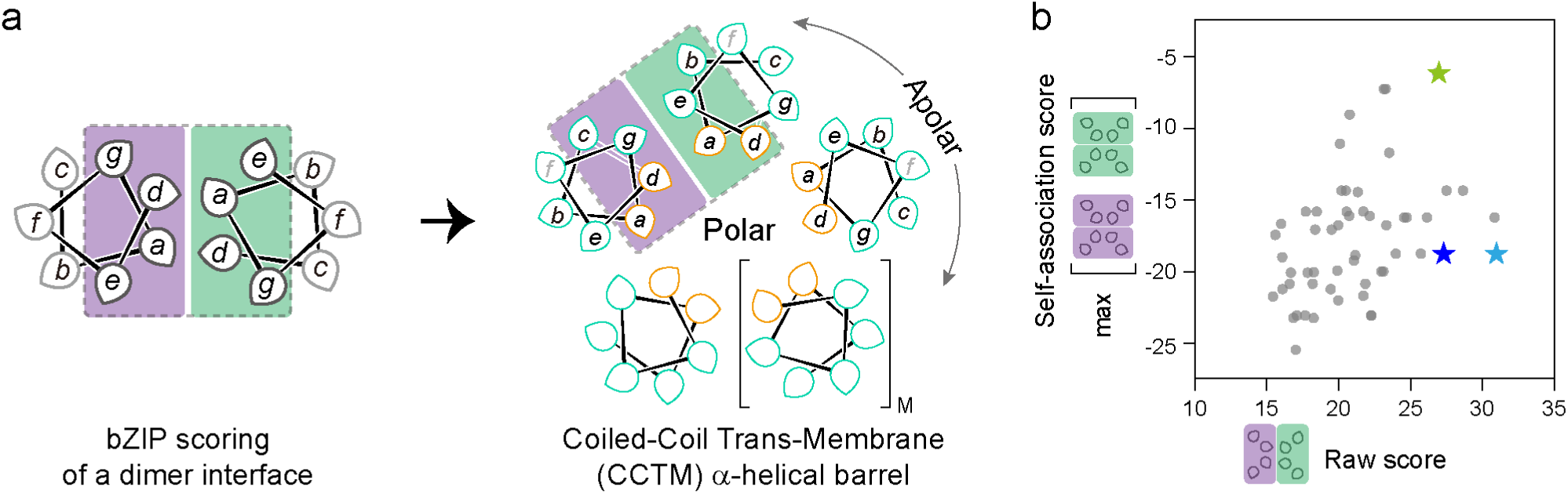
Design rationale for transmembrane coiled-coil barrels. (a) Design scheme of this work for obtaining transmembrane coiled-coil α-helical barrels through bZIP scoring.^31^ Helical wheels represent one of four heptad repeats in a peptide sequence. The pointed tips of the teardrop shapes indicate the orientation of side chains. The alphabetical ***a*** – ***g*** registers of coiled coils are shown in italics. Hydrophobic and polar residues are shown in cyan and orange, respectively. The design interface shown with purple and green squares is applied to bZIP scoring to obtain a Raw score. This design targets Type-II coiled-coil^3^ barrels with oligomer states ranging from 5 – 8 (M = 1 – 4). (b) bZIP scoring of design interfaces (Raw score) and alternative interfaces (Self-association score). Final fitness scores are calculated as the difference between Raw and Self-association scores. This gave CCTM-S_a_V_b_I_c_N_d,_ shown as a pale blue star, as the highest-scoring sequence. Pale green and blue stars represent CCTM-T_a_V_b_I_c_N_d_ and CCTM-S_a_V_b_A_c_N_d_, respectively.

We have previously exploited this ‘***gade***-based’ design strategy to generate water-soluble^11,29^ and membrane-spanning αHBs.^30^ For the computational design of water-soluble αHBs,^11^ we screened for mutually complementary pairs of ***g***-***a***-***d-e*** interfaces using bZip scoring^31^ as a proxy for coiled-coil packing. One hexameric barrel of these optimized soluble designs provided the basis for our subsequent rational design of a transmembrane αHB.^30^ Herein this work, to realize designs independent of template soluble counterparts, we combined these approaches, applying computational design and bZip scoring directly to deliver αHB ion channels with varied stoichiometries (Figure 1a). We name these designs Coiled-Coil Trans-Membrane (CCTM) αHBs. We further qualify this CCTM nomenclature with a suffix based on the designed sequence as follows.

We applied the following constraints to our CCTM designs: Residues at the ***b*** and ***c*** sites on the αHB exterior surface were constrained to be hydrophobic (A, I, L, V), as the barrel exterior must be membrane compatible. As the ***f*** position is not involved in helical interfaces, it was not specified during the scoring process. We limited interior-facing ***a*** and ***d*** positions to combinations of uncharged polar residues (N ,T, S) to define the ion-conducting channels. Gln was excluded since it gave lower Raw scores when included in sequences in our trial scorings (data not shown). Finally, a common feature of water-soluble αHBs is Ala at one or both of the ***e*** and ***g*** positions.^11, 29, 32^ Given the importance of these small residues at inter-helix contacts in membrane-spanning helices,^33, 34^ ***e*** and ***g*** positions were constrained to be Ala. As each peptide in a Type-II αHB requires two heterotypic packing seams (design interface) that share residues (Figure 1a), we calculated a fitness score for each of the 144 (4 x 3 x 3 x 4 for ***a – d*** positions) possible combinations of these heptad repeats. This was done by subtracting the score of the highest homo-typic interface (Self-association score) from that of the designed hetero-typic interface (Raw score). This scoring system aimed to maximize the specificity of helix-helix packing at the design interface. After filtering by the Raw score, this gave 56 hits. From these hits, ASVINA***f*** (in ***gabcdef*** register) had the highest fitness score (Figure 1b, Supplementary Table S1) and was chosen for further testing. A full sequence was generated with four heptads and ***f*** positions of W-L-L-W (Supplementary Table S2), commonly found at protein-membrane interfaces.^35^ We named this sequence CCTM-S_a_V_b_I_c_N_d_.

Following our previous designs,^11, 30^ atomistic models of parallel, blunt-ended, CCTM-S_a_V_b_I_c_N_d_ barrels were constructed. αHBs with oligomer states ranging from 4 to 8 were built using optimization of Crick parameterized α-helical coiled-coil oligomers within ISAMBARD.^36, 37^ These were assessed by empirical all-atom force field calculations using the Bristol University Docking Engine (BUDE).^38^ This gave an monotonically increasing model quality as oligomerization increased, with no clear local minima (Supplementary Table S3). We then assessed the stability of these oligomers in 1,2-diphytanoyl-*sn*-glycero-3-phosphocholine (DPhPC) lipid bilayers using all-atom molecular dynamics (MD) simulations. For all oligomers, the CCTM-S_a_V_b_I_c_N_d_ barrel structure equilibrated within the first 50 ns and was stable through the duration of the 210 ns simulation. (Supplementary Figure S1). Thus, unfortunately, neither energy minimization of parametrized αHBs nor MD simulations provided a definitive assessment of CCTM-S_a_V_b_I_c_N_d_ stoichiometry. Therefore, we turned to experimental characterization of the CCTM-S_a_V_b_I_c_N_d_ peptide.

### CCTM-S_a_V_b_I_c_N_d_ forms a pentameric membrane-spanning ion channel

A 36-residue version of CCTM-S_a_V_b_I_c_N_d_ was made by Fmoc solid-phase peptide synthesis, purified by reversed-phase HPLC and confirmed by MALDI-TOF mass spectrometry (Supplementary Figure S2 and S3). This included an *N*-terminal tetralysine tag and linker (KKKKGS) to enhance solubility, to aid purification, and to direct *C*-terminal insertion into the membrane (Supplementary Table S2).^2, 18^

The secondary structure of CCTM-S_a_V_b_I_c_N_d_ was assessed by circular dichroism (CD) spectroscopy. This indicated ≈75% helicity in phosphate-buffered saline (PBS) at pH 7.4 in the presence of either 0.35% pentaethylene glycol monooctyl ether (C8E5) or 1.5% n-octyl-β-D-glucoside (OG) (Supplementary Figure S4). This corresponds to the central four heptads (78%) of the sequence being helical. In the presence of 1,2-dimyristoyl-*sn*-glycero-3-phosphocholine (DMPC) large unilamellar vesicles, the high α-helical content was retained up to 85 °C, while the spectrum was distorted in buffer-only solution (Figure 2a). Analytical Ultracentrifugation (AUC) sedimentation-equilibrium experiments revealed CCTM-S_a_V_b_I_c_N_d_ to be pentameric in the presence of C8E5 detergent (Figure 2b).

**Figure 2.**
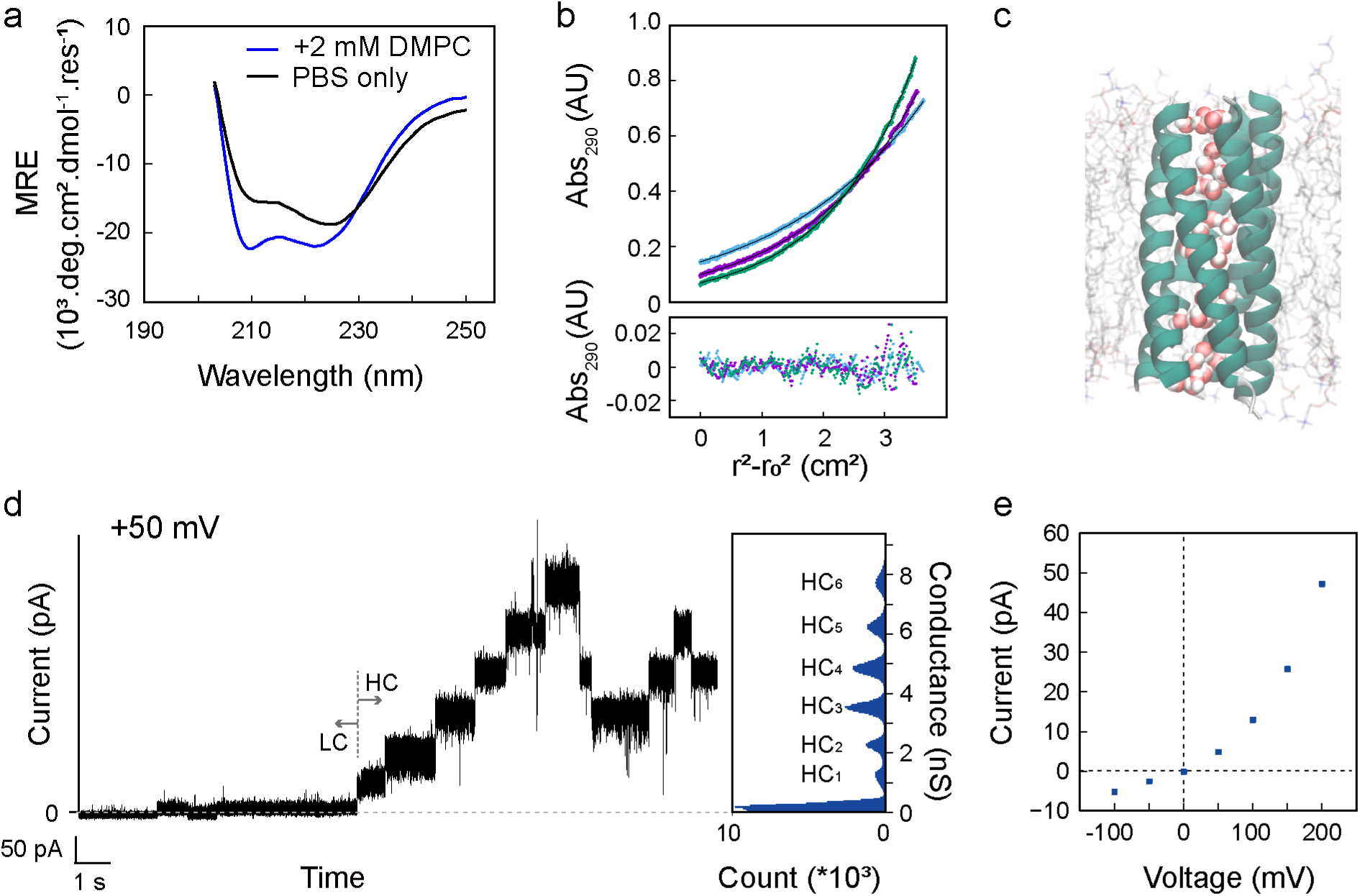
Biophysical and molecular dynamics (MD) characterizations of the CCTM-S_a_V_b_I_c_N_d_ peptide. (a) CD spectra of 20 μM CCTM-S_a_V_b_I_c_N_d_ in PBS measured at 85°C in the presence of large unilamellar vesicles formed by 2 mM dimyristoylphosphatidylcholine (DMPC) (blue line) and in the absence of vesicles (black line). (b) Analytical ultracentrifugation (AUC) plot; single-component fitting curve (top) and fitting residuals (bottom) from sedimentation-equilibrium data. Rotor speeds: 21 (blue), 24 (purple), and 27 (green) krpm. The estimated molecular weight of the assembly was 17925 Da (4.9 × monomer mass, 95% confidence limits 17772–18075 Da). (c) Snapshot after the 200 ns MD simulation of the CCTM-S_a_V_b_I_c_N_d_ pentamer in a 1,2-diphytanoyl-sn-glycero-3-phosphocholine (DPhPC) bilayer with water molecules (space-filled) in the central lumen. (d) Current recording and conductance histogram of CCTM-S_a_V_b_I_c_N_d_ (230 nM), low conductance (LC) and high conductance (HC) states at +50 mV. (e) Current-voltage plot of the LC state of CCTM-S_a_V_b_I_c_N_d_ (10 nM). For current recordings, the electrolyte was 1 M KCl in 10 mM Tris buffer (pH 8.0); signals were filtered at 2 kHz and sampled at 10 kHz.

Informed by these experiments, we took the pentameric model for CCTM-S_a_V_b_I_c_N_d_ into more-extensive MD simulations. From these, the barrel remained open, there was spontaneous permeation of water into the central channel, which then remained hydrated for the whole trajectory (Figure 2c, Supplementary Figure S5).

Next, we tested the channel-forming properties of CCTM-S_a_V_b_I_c_N_d_ using real-time single-channel electrical recordings in planar lipid bilayers. CCTM-S_a_V_b_I_c_N_d_ inserted into DPhPC bilayers and formed channels at positive applied voltages (Figure 2d); we observed an initial low-conductance (LC) state exhibiting ≈0.1 nS (230 nM peptide, + 50 mV, pH8, 1M KCl), followed by multiple distinct steps at higher conductance (HC_1,2,..n_, 1.6 – 1.9 nS). Using lower peptide concentrations (10 nM), it was possible to stabilize the LC at applied voltages up to +100 mV, and to characterize each HC step conductance (HC_1_ = 1.31 ± 0.04 nS, HC_2_ = 1.69 ± 0.09, HC_3_ = 1.78 ± 0.04, and HC_4_ = 1.88 ± 0.01 nS, at +200 mV). An I-V plot of the LC of CCTM-S_a_V_b_I_c_N_d_ was non-linear (Figure 2e). Transitions from LC to the HC states were voltage-dependent: application of >150 mV caused the channel to switch from LC to HC states; and lowering the applied voltage returned the system to the LC state (Supplementary Figure S6).

### CCTM channel stoichiometry can be altered by coiled-coil sequence and geometry

To test whether the properties of CCTMs could be modulated by rational changes in peptide sequence, we revisited the computational search for CCTM sequences. At the design interface, apart from fixed Ala at ***e*** and ***g*** positions, side chains at ***a***, and to a lesser extent at ***c***, point towards neighboring helices (Figure 1a). We reasoned that the relative orientations of the helical interfaces, which contribute to coiled-coil stoichiometry, might be sensitive to variations at these sites. Thus, we searched for single-or double-residue substitutions in the original ASVINA***f*** repeat at these positions with high-ranking design scores (Figure 1b, Supplementary Table S1). We selected ASVANA***f*** (CCTM-S_a_V_b_A_c_N_d_), with the smaller Ala at the ***c*** positions in place of Ile; and ATVINA***f*** (CCTM-T_a_V_b_I_c_N_d_) with the bulkier Thr at ***a*** positions instead of Ser. ATVINA***f*** gave a lower fitness score than ASVINA***f*** and ASVANA***f*** due to the relatively high self-association score of the I_c_N_d_A_g_T_a_ homo-typic interface (Supplementary Table S1). Nonetheless, since the self-association score was still negative and unfavorable, we chose ATVINA***f*** as a systematic modification to the parent CCTM-S_a_V_b_I_c_N_d_.

To probe the effect of membrane interfacing positions of the barrel-forming peptide sequences on channel formation, we also substituted the Trp at the ***f*** position of the first heptad of the parent CCTM-S_a_V_b_I_c_N_d_ with Lys to create CCTM-S_a_V_b_I_c_N_d_-KLLW. Like Trp, Lys is frequently found near the membrane surface region in natural membrane proteins,^39^ where its alkyl chain and positively charged terminal amino group are thought to “snorkel” at the membrane interface and compensate for hydrophobic mismatch between the lipid bilayer and peptide.^40, 41^ As a further probe of the impact of steric factors on helix-helix packing, we also sought to disrupt the interaction via chemical modification with iodoacetamide of single Ser→Cys substitution at residue 20 (S20C) in the parent CCTM-S_a_V_b_I_c_N_d_.

We synthesized the new CCTM peptides (Supplementary Table S2) and characterized them in C8E5 detergent solution by CD spectroscopy and AUC (results are summarized in Supplementary Figures S4 and S7, respectively). CD spectroscopy confirmed that CCTM-S_a_V_b_I_c_N_d_-S20C, CCTM-S_a_V_b_A_c_N_d_ and CCTM-T_a_V_b_I_c_N_d_ were all ≈75% helical, *i.e.,* similar to CCTM-S_a_V_b_I_c_N_d_. As expected, like the parent sequence, CCTM-S_a_V_b_I_c_N_d_-S20C gave a pentamer by AUC (17093 Da (4.7 × monomer mass), 95% confidence limits 16772 – 17373 Da). When the Cys residue was alkylated with iodoacetamide to yield CCTM-S_a_V_b_I_c_N_d_-S20C-ace, this formed a hexamer by AUC (22838 Da (6.1 × monomer mass), 95% confidence limits 22350 – 23317 Da) and retained its helical structure. This is consistent with the bulkier modified side chain at a central ***a*** pushing the helices apart from the inside to favor a higher oligomer state. For CCTM-S_a_V_b_A_c_N_d_ and CCTM-T_a_V_b_I_c_N_d_, AUC revealed that both were heptameric, returning molecular weights of 25969 Da (7.0 × monomer mass, 95% confidence limits 25529 – 26420 Da) and 24487 Da (7.0 × monomer mass, 95% confidence limits 24214 – 24761 Da), respectively. Again, these increases can be rationalized from the side-chain substitutions: the Thr@***a*** pushes the helices apart from the channel; the Ala@***c*** allows closer contacts between helices, and so both favorable barrels with more helices. CCTM-S_a_V_b_I_c_N_d_-KLLW was helical by CD spectroscopy (Supplementary Figures S4), but AUC indicated multiple oligomeric states (Supplementary Figures S7).

Based on these experimental oligomer states, we built structural models of the coiled-coil barrels (Figure 3). Heptameric CCTM-S_a_V_b_A_c_N_d_ and CCTM-T_a_V_b_I_c_N_d_ barrels were built and optimized in ISAMBARD. For CCTM-S_a_V_b_I_c_N_d_-S20C-ace, an optimized hexameric model of CCTM-S_a_V_b_I_c_N_d_ was altered to cysteine acetamide at S20C. Subsequent 210 ns all-atom simulations within a DPhPC bilayer indicated that like CCTM-S_a_V_b_I_c_N_d_ (Figure 3a), The CCTM-S_a_V_b_I_c_N_d_-S20C-ace also showed a circular barrel after the 210 ns simulation (Figure 3b, Supplementary Figure S8). The CCTM-S_a_V_b_A_c_N_d_ showed higher plasticity in helix-helix interface angles, resulting in an elliptical barrel (Figure 3c), while the CCTM-T_a_V_b_I_c_N_d_ was a circular but staggered barrel (Figure 3d, Supplementary Figure S9).

**Figure 3.**
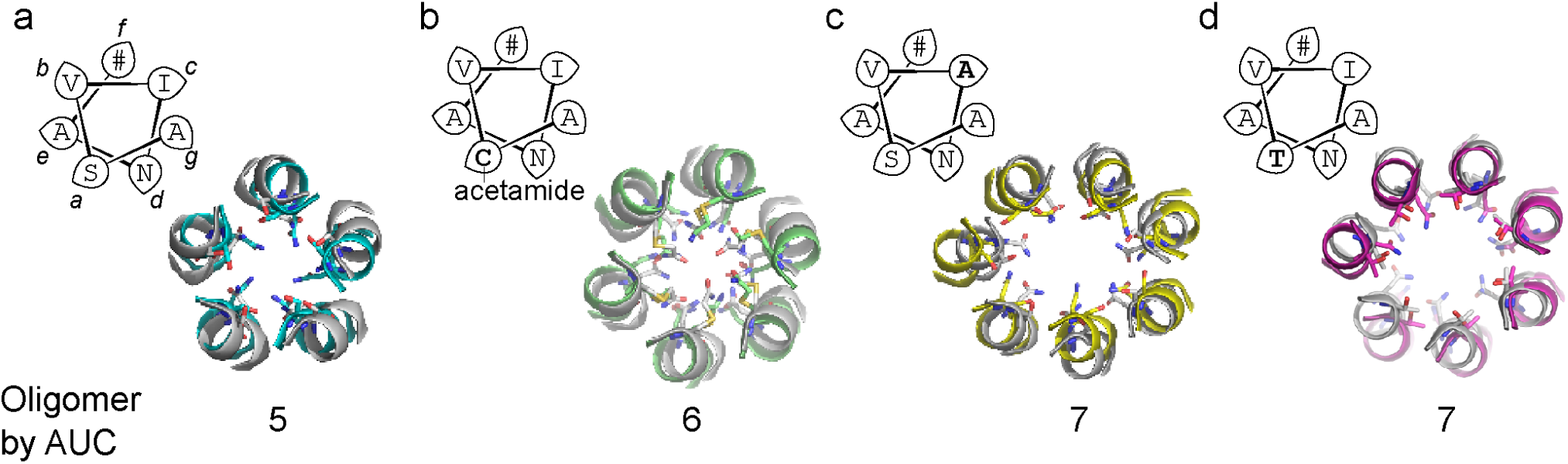
Varying CCTM stoichiometry. Helical wheels, model structures optimized for coiled coils (gray) and snapshots after 210 ns MD simulation (colored) as slice views at the third heptad, and experimental oligomeric states of designed peptide barrels by AUC, for the CCTM-S_a_V_b_I_c_N_d_ pentamer (a), CCTM-S_a_V_b_I_c_N_d_-S20C-ace hexamer (b), CCTM-S_a_V_b_A_c_N_d_ heptamer (c), and CCTM-T_a_V_b_I_c_N_d_ heptamer (d). Sidechains at ***a*** and ***d*** positions are shown as sticks. Oligomeric states were determined by analytical ultracentrifugation (AUC) using 20 – 25 μM peptide samples in PBS with 0.35% pentaethylene glycol monooctyl ether.

### CCTM channel function depends on coiled-coil geometry

Single-channel electrical recordings in planar lipid bilayers were used to characterize the CCTM variants. CCTM-S_a_V_b_A_c_N_d_, CCTM-T_a_V_b_I_c_N_d_, CCTM-S_a_V_b_I_c_N_d_-S20C-ace, and CCTM-S_a_V_b_I_c_N_d_-KLLW all inserted into bilayers and formed channels. As with the parent peptide, for all of these, we observed a low conductance state (LC), followed by multiple distinct steps at higher conductance (HC_1,2,..n_) (Figure 4a–d). Like CCTM-S_a_V_b_I_c_N_d_, the transition between LC and HC states was voltage-dependent, and particularly CCTM-S_a_V_b_I_c_N_d_-KLLW showed stable channel formation (Supplementary Figure S10). Surprisingly, we observed little variation in the incremental conductances for the HC state, regardless of the modifications to the CCTM sequence: HC_1_ for CCTM-T_a_V_b_I_c_N_d_, CCTM-S_a_V_b_A_c_N_d_, CCTM-S_a_V_b_I_c_N_d_-S20C-ace, and CCTM-S_a_V_b_I_c_N_d_-KLLW were 0.73 ± 0.02 (+200 mV), 0.54 ± 0.07 (+200 mV), 0.68 ± 0.1 (+150 mV), 0.78 ± 0.05 nS (+150 mV), respectively, followed by HC_2,..n_ 1.0 – 2.1 nS (Figure 4e). In contrast, for all CCTMs, we observe a marked variation in the magnitude of the LC state. The LC of CCTM-S_a_V_b_A_c_N_d_ (0.038 ± 0.010 nS) was the smallest, followed by CCTM-S_a_V_b_I_c_N_d_-S20C-ace (0.059 ± 0.023 nS), CCTM-T_a_V_b_I_c_N_d_ (0.062 ± 0.012 nS), CCTM-S_a_V_b_I_c_N_d_ (0.089 ± 0.024) and CCTM-S_a_V_b_I_c_N_d_-KLLW (0.111 ± 0.017 nS) (N=25), respectively (Figure 4f).

**Figure 4.**
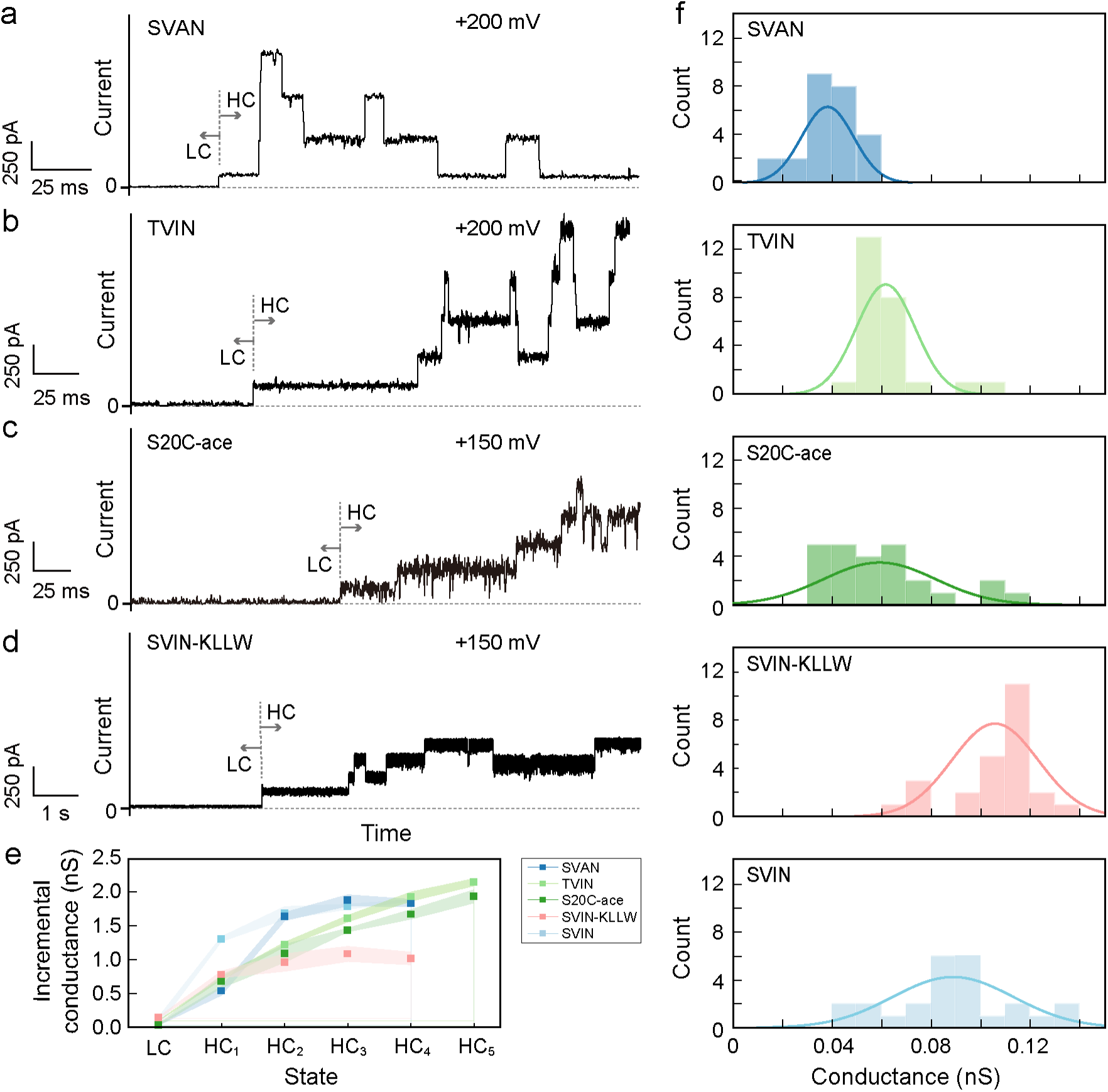
Electrical conductance properties of the designed peptides. (a-d) Current recording of CCTM-S_a_V_b_A_c_N_d_ at +200 mV (a), CCTM-T_a_V_b_I_c_N_d_ at +200 mV (b), CCTM-S_a_V_b_I_c_N_d_-S20C-ace at +150 mV (c), and CCTM-S_a_V_b_I_c_N_d_-KLLW at +150 mV (d), exhibiting low-conductance (LC) and high-conductance (HC) states. For current recordings, peptide concentrations were 10–50 nM; the electrolyte was 1 M KCl in 10 mM Tris buffer (pH 8.0); signals were filtered at 2 kHz and sampled at 10 kHz. (e) Increment conductances HC_n_ between *n*th and (*n-1*)th steps of the HC state CCTM-S_a_V_b_I_c_N_d_ (pale blue), CCTM-S_a_V_b_A_c_N_d_ (blue), CCTM-T_a_V_b_I_c_N_d_ (pale green), CCTM-S_a_V_b_I_c_N_d_-S20C-ace (green) and CCTM-S_a_V_b_I_c_N_d_-KLLW (pink), respectively. The error area represents the standard deviations of three experimental repeats. (f) Histograms and fitted curves to a Gaussian function of unitary conductance of the LC state (N=25) measured at 50 – 200 mV. The color scheme is the same as panel (e) for all peptides.

### Large CCTM current steps indicate a single pore

It was not clear whether each step of the HC state was produced by expansion and contraction of a single pore, or arose from multiple barrels inserting into the bilayer. Therefore, we probed the HC state in more detail through optical single-conductance recordings (oSCR) with droplet interface bilayers (DIB) (Figure 5a).^42, 43^ This allowed us to monitor the conductance of the ensemble of channels in a bilayer, as well as the location and mobility of the ion flux through individual channels. In oSCR, the transmembrane flux of Ca^2+^ ions are visualized by TIRF microscopy using Fluo-8H, a Ca^2+^-sensitive dye.

**Figure 5.**
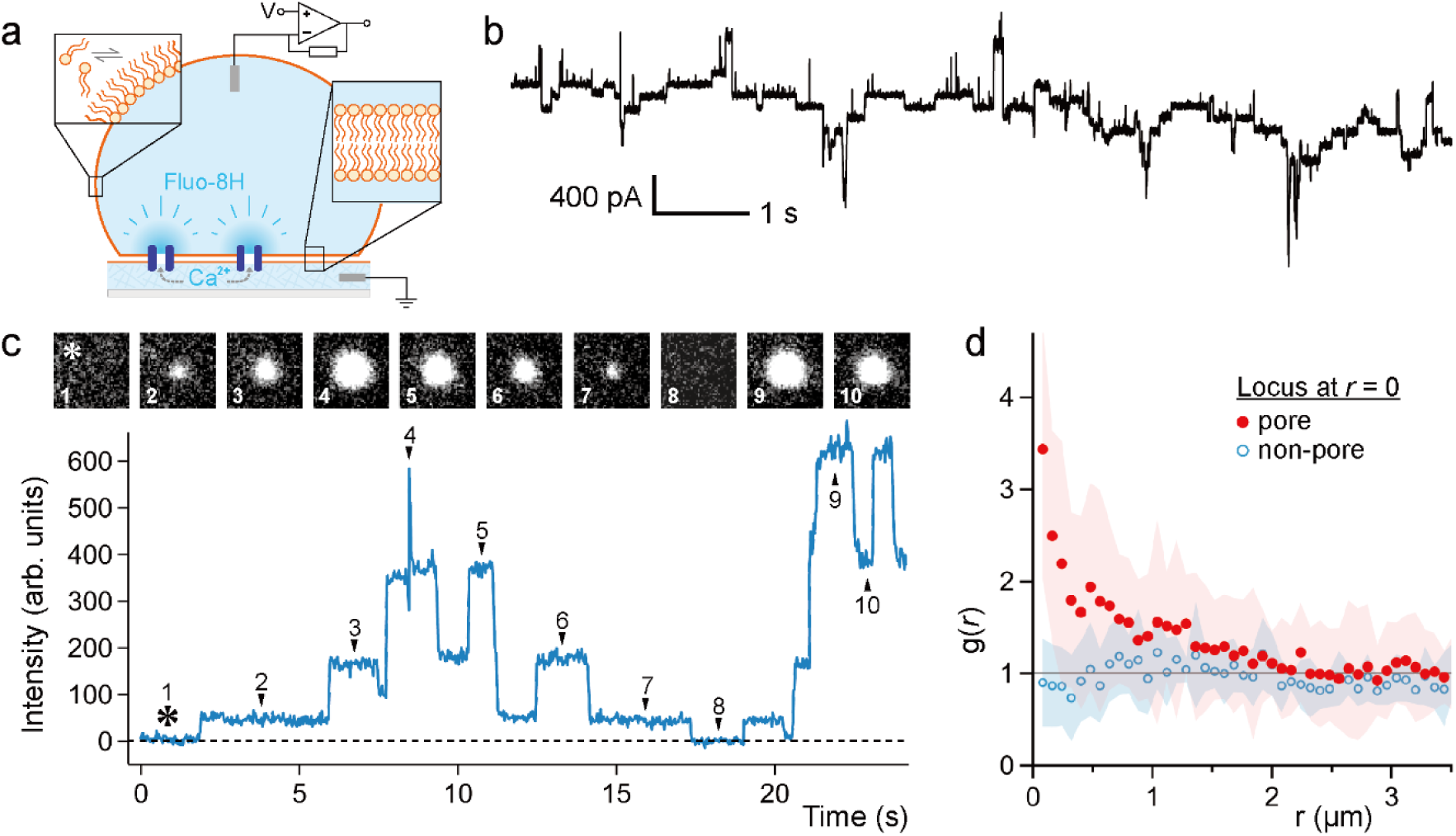
CCTM-S_a_V_b_I_c_N_d_ [KLLW] in droplet interface bilayers. (a) Diagram of the optical single-channel recording (oSCR) setup. Lipid monolayers (orange) spontaneously assemble on a planar hydrogel layer spun onto a coverslip (light grey) and an aqueous droplet in the presence of lipid-in-oil. When the droplet contacts the hydrogel, a bilayer is formed with Ca^2+^ ions on one side and the Ca^2+^-sensitive dye Fluo-8H on the other. Pores formed in the membrane (by CCTM-S_a_V_b_I_c_N_d_ [KLLW], purple) allow the passage of Ca^2+^ across the membrane, where they bind with Fluo-8H. The plume of Ca^2+^-bound dye emerging from the site of the pore (dark blue) was imaged using total internal reflection fluorescence microscopy. Current recordings were made through electrodes in the droplet and the hydrogel. (b) Multiple conductance steps from 400 nM CCTM-S_a_V_b_I_c_N_d_ [KLLW] in 1,2-diphytanoyl-sn-glycero-3-phosphocholine at -90 mV. (c) oSCR showing that a single, isolated pore fluctuates between multiple conductance states. The bilayer was held at -60 mV, and the peptide concentration was 400 nM. Images above the trace show the oSCR signal varying in size.

CCTM-S_a_V_b_I_c_N_d_ [KLLW] was inserted into DIBs made from DPhPC and recordings showed the HC state (Figure 5b), as in the measurements with planar lipid bilayers. Under TIRF illumination, bright puncta of discretely varying intensity were observed on the surface of the bilayer, indicating transmembrane pores that conduct Ca^2+^. When single pores were isolated, oSCR signal intensity fluctuated in discrete, multiple levels (Figure 5c). Further experiments were performed with a small fraction (0.5 mol%) of CCTM-S_a_V_b_I_c_N_d_ [KLLW], *C*-terminally labeled with Cy5 dye. This allowed individual CCTM monomers to be observed diffusing as single molecules on the membrane surface. We then correlated the positions of the monomers with the channels observed using oSCR. A radial distribution function (RDF, g(r)) indicates the probability of finding peptides in the region of the pore (r = 0). Figure 5d shows the RDF computed around a single channel and at arbitrary locations on the bilayer surface (not at the pore site). As expected for freely diffusing molecules, g(r) was unity where there were no channels (blue line); i.e., there was an equal probability of finding a labeled peptide in all locations. However, near the channel locus (red line), g(r) > 1 indicating an increased probability of finding peptides near the pore (near r = 0). Moving away from the locus, g(r) dropped to 1 again. Together, these data indicate that the HC states are single channels of the designed peptides but that they change dynamically in radius in discrete steps. We interpret this as stochastic addition and loss of peptides from the transmembrane barrel.

To test if even larger stoichiometries than the designed barrels are possible, we performed 1 μs all-atom MD simulations of 10-mer, 15-mer, and 20-mer CCTM-S_a_V_b_I_c_N_d_ barrels in DPhPC bilayers at 353 K in the absence and presence of membrane potentials (Supplementary Figure S11). All barrels showed collapsed structures in the absence of the membrane potential. In contrast, at +200 mV membrane potential the 10-mer maintained the elliptical barrel in the duration of the simulation. The 15-mer and 20-mer barrels exhibited more distorted channels at +200 mV, but again did not collapse. The conductance of the 10-mer elliptical barrel was estimated to be ≈1.2 nS (1M KCl, +200 mV, 300 K) from the MD trajectory (The raw estimate 3.6 ± 0.1 nS was divided by three with consideration of the three times faster diffusion of TIP3P water than physical water),^44^ which is in close proximity to the experimental conductance of HC_1_ (1.4 nS) of CCTM-S_a_V_b_I_c_N_d_. These results also indicate that at the HC state with high applied voltages CCTM peptides can form the more dynamic and larger single channels than the LC state.

The sequence starts with the membrane intact (*); numbers correspond to the points marked by arrowheads in the trace. Images represent ≈ 28 × 28 μm. (d) Mean radial distribution functions (RDFs) around CCTM-S_a_V_b_I_c_N_d_ [KLLW] pores (red, N = 10) and arbitrary points elsewhere on the membrane (blue, n = 8). Shaded regions indicate the standard deviation of the mean.

## Discussion

We have shown that the computational design of coiled-coil peptide assemblies can deliver a series of membrane-spanning α-helical barrels (αHBs) with ion-channel activity. Initially, we made two pentamers, one hexamer, and two heptamers. The parent design, CCTM-S_a_V_b_I_c_N_d,_ with the coiled-coil heptad repeat ***gabcdef*** = ASVINAx, forms a pentamer in the presence of detergent. Rationally designed variants of this show that the stoichiometry of the peptide can be altered. For instance, a Ser→Cys variant at one of four central ***a*** positions, and with the Cys modified by a bulky acetamide group (CCTM-S_a_V_b_I_c_N_d_-S20C-ace), forms a hexamer. Whereas, replacing all Ile at ***c*** with Ala results in a heptamer (CCTM-S_a_V_b_A_c_N_d_); and substituting all of the Ser at ***a*** with Thr gives another heptamer (CCTM-T_a_V_b_I_c_N_d_). In contrast to these observations, for designed water-soluble αHBs, the substitution at ***a*** with beta-branched side chains (e.g. Leu to Ile) results in smaller oligomer states,^29^ and ***c*** positions are normally simply made extended charged residues to promote interhelical salt bridges. This highlights challenges in modeling transmembrane peptide assemblies and predicting their stoichiometries without explicit consideration of membrane environments.^45^

CCTM-S_a_V_b_I_c_N_d_ and other variants form pores in a planar lipid bilayer, with stable low-conductance (LC) states at low potentials, and a high-conductance (HC) state at higher potentials. The conductance range of the LC state is consistent with a Type-II αHB model,^46^ in which sequence changes lead to changes in oligomeric state, which, in turn, alter the channel geometry and conductances. However, the HC states, which are more similar across the designs, exhibit discrete steps that could be due to either the insertion of additional pores or the recruitment/expulsion of peptides to alter pore size. To distinguish between these models, we conducted optical single-channel recordings (oSCR) with droplet interface bilayers (DIB). From these, we conclude that HC states arise from dynamic single pores that recruit and expel peptides. Molecular-dynamics (MD) simulations of the large barrels support this model as we observe collapse and fragmentation in the absence of membrane potential and stabilization of the barrel in the presence of the positive potential. Furthermore, the HC state of the designed peptides shows similar behavior to alamethicin, a natural antimicrobial peptide that forms barrel-stave pores in lipid bilayers and exhibits discrete changes in pore size.^47, 48^ The mechanism of the discrete current steps of alamethicin has been described as changes in stoichiometry by recruiting/releasing monomeric peptides corresponding to each step.^49–53^ The fact that the HC states of CCTM variants are essentially similar to alamethicin suggests a pore-formation mechanism (at least for this HC state) that is more general and less dependent on changes in sequence and specific helix-helix interaction; *i.e*, it is a general property of amphipathic membrane-spanning channel-forming helices like alamethicin and the CCTM designs.

By contrast, the conductance of the LC channels correlates with the location and sidechain bulkiness of the rational design modifications. However, notably, the stoichiometry and conductance here are not necessarily proportional. The electrical recordings of heptameric CCTM-S_a_V_b_A_c_N_d_ showed that Ala at ***c*** and ***g*** results in the smallest conductance among all variants. Based on the MD simulation, this is likely because the higher plasticity derived from small residues leads to the narrower elliptical channel. The conductances of heptameric CCTM-T_a_V_b_I_c_N_d_ and hexameric CCTM-S_a_V_b_I_c_N_d_-S20C-ace are greater than CCTM-S_a_V_b_A_c_N_d_ since it has a circular channel, but smaller than pentameric CCTM-S_a_V_b_I_c_N_d_ due to larger sidechains than Ser at the lumen. Furthermore, the similar LCs of CCTM-S_a_V_b_I_c_N_d_-KLLW and CCTM-S_a_V_b_I_c_N_d_ indicates that modifications at the barrel exterior have a limited impact on the conductance. Our conductance prediction of CCTM-S_a_V_b_I_c_N_d_, CCTM-T_a_V_b_I_c_N_d_, and CCTM-S_a_V_b_A_c_N_d_ barrel models with a series of oligomer states using HOLE program^54^ also showed that the conductance of pentamer to heptamer do not linearly increase in all peptides (Supplementary Figure S12). This would be because the subtle difference of the side-chain arrangement at the lumen governs the function for these small channels. Thus, our results deliver the non-trivial, sequence-stoichiometry-function relationships of membrane-spanning coiled-coil αHBs, which will provide a guide for tuning peptide channel functions.

Recent studies of redesigned αHBs have shown that van der Waals interactions between apolar side chains are crucial in stabilizing the helix–helix association of phospholamban-like pentameric peptides in a hydrophobic environment.^20^ Increasing the interaction surface area between monomers could also provide highly stable structures and help address the challenge in high-resolution structural analyses. Indeed, computational design of TMH4C4 has been shown to achieve this, resulting in a cryo-EM structure of the tetramer of long four-transmembrane helix subunits.^3^ This architecture is built from a water-soluble αHB with a hydrogen bonding network inside the pore, followed by apolar conversion of residues at the barrel exterior to maintain both tight helix–helix interactions and transmembrane topology. In our previous design of a peptide-based ion channel, we applied this strategy to a single-layer coiled-coil peptide barrel^2^ and found that it is essential to optimize both hydrogen bonding and steric packing, which had been proposed previously in designing membrane α-helix assemblies.^55^ Building on this, the work presented here demonstrates that computational design can successfully find packing solutions that employ polar and apolar side chains to stabilize αHBs and to introduce conductive channels. This has delivered the CCTM designs. The obtained stable packing enables biophysical characterization of the designed peptides, which we suggest represents the low-conductance states. Interestingly, however, high membrane potentials promote the recruitment (and expulsion) of additional peptide and the formation of dynamic channels of variable size. This highlights opportunities and challenges in designing membrane-spanning channel-forming peptides and proteins. Specifically, the *de novo* peptides that we describe exhibit dynamic structural and associated functional (conductance) changes, which has exciting potential for generating voltage-sensitive and other responsive channels; however, we will have to learn how to control this behavior and design it predictably to realize such systems. One route might be to exploit recent advances in computational design that takes into account conformational flexibility^56^ and that use peptide assemblies as rational seeds in the design of defined single-chain αHB proteins.^57–59^

## Materials and Methods

### Peptide design and modeling

Following our previous methodology,^11^ we used bZIP scoring^31^ to identify complementary pairs of coiled-coil interfaces of αHBs. The bZIP algorithm assigns scores to pairs of dimer interfaces defined by their ***g, a, d,*** and ***e*** (***gade***) residues of a heptad repeat with the alphabetical register (***abcdefg***). For two ***gade*** configurations to be compatible with a type-II coiled-coil interface,^60^ they must share two residues, such that the ***a*** and ***e*** residues of the first ***gade*** are the same as the ***g*** and ***d*** residues of the second. Pairings of ***gade*** configurations meeting this criterion and adhering to the residue choice rationale described in the Results section were screened with bZIP to identify those predicted to form heterotypic pairs. Each pair of ***gade*** configurations was assigned a fitness score, calculated by taking the Raw score for the desired ***gade***1 + ***gade***2 combination and subtracting from it the highest competing Self-association score (either ***gade***1 with ***gade***1 or ***gade***2 with ***gade***2). This score gives an indication of the degree to which the two interfaces are prone to form heterotypic combinations.

Atomistic models of the resulting CCTM barrels were built using the ISAMBARD framework^36^ for protein design. For each model, radius, pitch, and relative rotation of the helices were optimized using a differential evolution algorithm.^37^

### Molecular dynamics (MD)

Conventional all-atom MD simulations were performed for CCTM-S_a_V_b_I_c_N_d_, (5, 6, 7, 8, 10, 15, 20-mer), CCTM-T_a_V_b_I_c_N_d_ (heptamer), CCTM-S_a_V_b_*A*_c_N_d_ (heptamer), CCTM-S_a_V_b_I_c_N_d_-S20C-ace (hexamer), in a 1,2-diphytanoyl-*sn*-glycero-3-phosphocholine (DPhPC) lipid bilayer. Initial barrel structures were built using ISAMBARD,^36^ with a tetralysine tag (KKKKGS) attached to the N-terminus of each peptide in PyMOL (Version 2.5.2, Schrödinger, LLC). For the initial configuration of each simulation, the peptide barrel structure was placed at the center of the XY plane, and the Z coordinate was predicted by the PPM webserver.^61^ The structure was embedded in a DPhPC bilayer using Membrane Builder in CHARMM-GUI.^62, 63^ All Lys residues were protonated, consistent with a pH of 7.0. N-and C-termini of the peptides were acetylated and amidated, respectively. Each system was solvated and neutralized with 1 M KCl solution with a water thickness of 20 Å. The system size information is summarized in Supplementary Table S4.

All simulations were performed with versions 2.0 beta and 2.1 of the GENESIS MD program.^64–67^ The CHARMM36 protein force field,^68, 69^ SwissSidechain,^70^ CHARMM36 lipid force field,^71^ and TIP3P model^72^ were used for peptides, cysteine-S-acetamide residues, lipids, and water, respectively. Bonds with hydrogen atoms were constrained using SHAKE/RATTLE^73, 74^ for non-aqueous molecules and SETTLE^75^ for water. Long-range electrostatic interactions were evaluated by particle-mesh Ewald summation.^76, 77^ A cut-off distance of 12 Å for non-bonded interactions was used with an force-based switching function effective from 10.0 Å. The pairlist distance for non-bonded interactions was set to 13.5 Å. The system was energy-minimized and equilibrated following the procedure provided by CHARMM-GUI (summarized in Supplementary Table S5). Production simulations were performed at 300 or 353 K and 1 atm with isothermal-isobaric conditions (NPT ensemble) controlled by Bussi’s stochastic velocity rescaling thermostat and MTK barostat with semi-isotropic condition.^78^ A multiple time step integration using r-RESPA was employed,^79, 80^ with 3.5/7.0 fs and hydrogen mass repartitioning.^65^ In simulations with a membrane potential, a uniform electric field (E_z_) was applied to all atoms along the z-axis. The electrical potential ranges from *V = 0* to *V = L_z_E_z_*, where L_z_ is the length of the simulation box along the z-axis. For conductance estimates, we performed 100 ns MD under the NPT condition, after 10 ns equilibration at 300K from the last snapshot of the 1-μs simulation at 353K. Ion currents of the peptide pores were analyzed through computing instantaneous ion currents as described by Aksimentiev *et al*.^44^ MD simulations were performed on HOKUSAI-BW and FUGAKU in RIKEN, Japan. VMD^81^ and PyMOL 2.5.2 (Shrodinger, LLC) were used for visualization. Analyses of MD trajectories were performed on GENESIS and VMD.

### Peptides

Peptides were synthesized by Fmoc solid-phase synthesis with a CEM Liberty Blue automated instrument with in-line UV monitoring. Boc-Ser(Fmoc-Ala)-OH or Boc-Thr(Fmoc-Ala)-OH were used at the A-S/T position in the third heptad. The rest of the peptide after the dipeptide block was synthesized without microwave heating. Cleavage from the solid support was carried out with a mixture of trifluoroacetic acid (TFA)/triisopropylsilane/water (90/5/5, v/v/v). An additional 5% ethanedithiol was added to the cleavage mixture for all Cys-containing peptides. Peptides were purified by reversed-phase high-pressure liquid chromatography (HPLC) on a Vydac® TP C18 column (10 µm particle, 22 × 250 mm), followed by a second fractionation on a Phenomenex® Luna C8 column (5 μm particle, 10 mm × 250 mm). Fractions containing pure peptides were identified by analytical HPLC and MALDI-TOF mass spectrometry (MS) on a Bruker Ultraflex instrument (Supplementary Figure S3 and S13). The purified O-acyl peptides were subsequently rearranged to N-acyl peptides by treatment with ammonium bicarbonate (Supplementary Figure S2). See Supplementary Methods for more detailed information.

### Alkylation of CCTM-S_a_V_b_I_c_N_d_-S20C

An aqueous solution of iodoacetamide total alkylating agent (100 mM) was added to CCTM-S_a_V_b_I_c_N_d_-S20C (50 μM) with tris(2-carboxyethyl)phosphine hydrochloride (TCEP-HCl, 0.5 mM) in phosphate-buffered saline (PBS; 8.2 mM sodium phosphate, 1.8 mM potassium phosphate, 137 mM sodium chloride, 2.7 mM potassium chloride at pH 7.4)/TFE = 1/1 (v/v) in the dark. The reaction was monitored by analytical HPLC (Supplementary Figure S14) for 2 h and quenched with an aqueous solution of 1 M formic acid. The reaction product was purified on a Phenomenex® Kinetex C4 (100 × 4.5 mm) column with water + TFA (0.1%, buffer A) and MeCN + TFA (0.1%, buffer B), with a gradient of 40 to 95% buffer B over 30 min. The purified CCTM-S_a_V_b_I_c_N_d_-S20C-ace was identified with MALDI-TOF MS/MS (Supplementary Figure S15).

### Preparation of Cy5-labeled CCTM-S_a_V_b_I_c_N_d_

A Cys-Gly-CCTM-S_a_V_b_I_c_N_d_ peptide was synthesized, cleaved, and purified by reversed-phase HPLC and a Cy5 maleimide dye coupled to the Cys residue of the purified peptide. To achieve this, we added Cys-Gly-CCTM-S_a_V_b_I_c_N_d_ [KLLW] (0.05 μmol) in 500 μL of PBS (pH 6.6) to TCEP (1.25 mg, 0.5 μmol), followed by sulpho-Cy5 maleimide (Lumiprobe, Germany) in dimethyl sulfoxide (0.402 mg, 0.5 μmol, 10 mg ml^-1^); the mixture was rocked overnight. The dye-labeled peptide was purified by reversed-phase HPLC using a Phenomenex Luna C8 column (5 μm, 100 Å, 10 mm ID × 250 mm L) with water + TFA (0.1%, buffer A) and MeCN + TFA (0.1%, buffer B), with a gradient of 40 to 100% buffer B over 30 min. The purified Cy5-labeled CCTM-S_a_V_b_I_c_N_d_ [KLLW] was identified using analytical HPLC and MALDI-TOF MS (Supplementary Figure S16).

### Circular dichroism (CD)

CD spectra were measured using JASCO J-810, J-815 (Jasco, UK) or J-1500 (Jasco, Japan) spectropolarimeters with Peltier temperature controllers. Peptide samples with detergent micelles were prepared as 20 μM solutions in PBS with 1.5% n-octyl-β-D-glucoside (OG), or 0.35% pentaethylene glycol monooctyl ether (C8E5). For samples with lipid vesicles, large unilamellar vesicles were prepared prior to peptide addition. A chloroform solution of 1,2-dimyristoyl-sn-glycero-3-phosphocholine (DMPC, final 2 mM) was dried under nitrogen gas followed by under vacuum overnight. The obtained lipid film was hydrated with PBS at 40°C for 30 min and vortexed. The suspension was subjected to five freeze/thaw cycles and subsequently extruded 35 times through a 100-nm polycarbonate membrane at 40°C by using Avanti Extruder Kit (Avanti Polar Lipids). The obtained LUV suspension was mixed with a peptide solution in PBS (peptide to lipid molar ratio: 1:100). Spectra were accumulated using a 1-mm path length quartz cuvette at 20°C unless specified. For each dataset (in deg), baselines from the same buffer and cuvette were subtracted, and data were normalized for amide bond concentration and path length to yield mean residue ellipticity (deg cm^2^ dmol^-1^ res^-1^).

### Analytical ultracentrifugation (AUC)

Sedimentation equilibrium experiments were performed with a Beckman Optima XL-I or XL-A analytical ultracentrifuge using An-50 Ti or An-60 Ti rotors (Beckman Coulter). Samples were prepared in PBS containing 0.35% C8E5 at peptide concentrations of 20–25 μM and centrifuged at 21–27 krpm. Datasets were fitted to single-component models using Ultrascan II (http://www.ultrascan.uthscsa.edu). The partial specific volume for each peptide and the buffer density were calculated using Sednterp (http://sednterp.unh.edu/).

### Single-channel recordings

Membrane insertion and channel formation of CCTM peptides were studied in planar lipid bilayers under applied potentials. Bilayers were formed using DPhPC (Avanti Polar Lipids) by employing the classic Montal and Mueller technique.^82^ A 25-μm Teflon film (PTFE) partition, with an aperture of ∼60 μm in diameter and painted with 1% hexadecane, was placed between *cis* and *trans* compartments of a flow cell. Both compartments were filled with the electrolyte solution (1 M KCl, 10 mM Tris-HCl, pH 8.0). DPhPC in *n*-pentane (5 μL, 5 mg/mL) was added to both sides to allow membrane formation over the aperture by lowering the buffer level below the aperture and raising it. Standard Ag/AgCl electrodes with 3 M KCl/agar bridges were placed in each chamber to measure the ion current. For single-channel measurements, unless otherwise specified, 10 nM peptide was added to the cis side of the chamber for insertion into the membrane under an applied potential. The *cis* side was connected to the ground electrode. Spontaneous peptide insertion was obtained under an applied voltage of +50 mV (230 nM peptide) or +200 mV (10 nM peptide). Current was amplified using an Axopatch 200B amplifier, digitized with a Digidata 1440A A/D converter, and recorded using pClamp 10.3 acquisition software (Molecular Devices, San Jose, CA, USA) with a low-pass filter frequency of 2 kHz and a sampling frequency of 10 kHz. Data were analyzed and prepared for presentation with pClamp (version 10.3, Molecular Devices) and Origin 9.0 or 2022b.

### Electrical and optical recordings with droplet interface bilayers (DIBs)

DIBs were prepared as described previously.^42^ Briefly, a 0.75% (wt/vol) ultralow temperature gelling agarose (Sigma-Aldrich) solution in Milli-Q water was prepared and heated to 90°C; then 140 μL was spun onto a coverslip (Menzel-Gläzer, ThermoFisher Scientific). The coverslip was affixed to the underside of a custom-made poly(methyl methacrylate) device featuring 16 wells, 1 mm in diameter. A 1% (wt/vol) agarose solution containing 1 M KCl (or 500 mM CaCl_2_ for optical SC recording [oSCR] experiments), buffered with 25 mM Tris-HCl at pH 8.0, was flowed into the device, where it made contact with and hydrated the substrate agarose.^42^ The wells were filled with hexadecane containing DPhPC at 8.7 mg/mL and incubated for 1 h to allow for monolayer formation on the substrate. Meanwhile, aqueous droplets (∼100 nL) were incubated in the same lipid-in-oil solution for additional monolayer formation. The droplets contained 1 M KCl, 25 mM Tris-HCl, 375 μM EDTA, and 23 μM Fluo-8H, pH 8.0, in addition to unlabeled (400 nM) and Cy5-labeled CCTM-S_a_V_b_I_c_N_d_ [KLLW] (2 nM), as required. After incubation, droplets were pipetted into the wells, where they sedimented onto the substrate, forming bilayers by the contact of the droplet with agarose-associated monolayers.

Ag/AgCl electrodes were inserted into the hydrating agarose and electrical access was granted to the top of the droplet using a micromanipulator. Voltages were applied and bilayer currents were recorded using an Axopatch 200B patch-clamp amplifier and headstage (Axon Instruments, Molecular Devices, California, USA), with signals recorded and filtered at 1 kHz. Devices were placed within a Faraday cage atop an inverted microscope (TiE Eclipse, Nikon, UK). Fluorescence imaging was carried out using a 60× total internal reflection fluorescence (TIRF) objective lens (oil immersion, 1.45 NA; Nikon). Excitation of the bilayers and collection of fluorescent signals were carried out using the same objective. For excitation of Ca^2+^-bound Fluo-8H, a 473-nm continuous-wave laser beam was used, with 1–4 mW of power at the back focal plane of the objective (Vortran Laser Technology, Sacramento, California, USA); fluorescence and excitation signals (λ_Fluo-8H emission max._ = 514 nm) were separated by a dichroic mirror (ZT 488/640; Chroma, Vermont, USA); and the collected signal from the bilayer was transmitted through a Brightline bandpass 550/88-nm emission filter (Semrock, Rochester, New York, USA) to eliminate stray excitation wavelengths. Cy5 was excited using a 644-nm continuous-wave laser beam, with 4–7 mW of power at the back focal plane of the objective (Vortran Laser Technology). Emitted fluorescence was passed through a 664-nm longpass dichroic mirror and 680/42-nm bandpass emission filter (Semrock). Images were recorded at 33 Hz using an electron-multiplying charged-coupled device camera (iXon+; Andor Technology, Belfast, UK). All experiments were conducted at room temperature (approximately 21°C). For calculating radial distribution functions (RDFs), g(r), after DIB formation, a region of the bilayer containing channels was imaged over several hundred frames. This area was simultaneously illuminated by blue and red lasers to excite Fluo-8H and Cy5, respectively, with the emitted light spectrally separated by a single camera. After spatially registering this two-color fluorescence, the 2D Gaussian fitting of the oSCR signal allowed sub-pixel localization of the CCTM channels. A tracking algorithm was used to locate Cy5-labeled peptides in the red channel on the same membrane patch, regardless of whether they were part of a transmembrane pore. Then RDF were calculated for labeled peptides using the pore centers (as determined by 2D Gaussian fitting of the Ca^2+^-flux images) as r = 0. g(r) is a normalized count of the number of particles (in this case, peptides) found at a given distance r from a reference point (in this case, the pore center) within a circular shell of thickness dr; normalization is by the area of the shell and for the particle density, such that g(r)→1 as r→∞. In this way, the RDF was calculated every 80 nm from the reference point, with a shell thickness dr of 80 nm. This value is approximately equal to the localization precision for single molecules.

Electrical data were recorded using WinEDR electrophysiology software (John Dempster, Strathclyde University, UK). Image analyses were carried out using Fiji image-processing software,^83^ and single-particle tracking was performed using the TrackMate plugin, using its Laplacian of Gaussian algorithm.^84^ Numerical data and RDF analyses were performed in Igor Pro (Wavemetrics, Oregon, USA) using custom-written procedures.

## Acknowledgements

We are grateful to Dr. Aimee Boyle for the insightful discussion and advice. We thank the RIKEN Molecular Structure Characterization Unit for CD measurement and the CBS Support Unit for Bio-Material Analysis for peptide synthesis. We acknowledge the computing time granted by the RIKEN Center for Computational Science (HOKUSAI BigWaterfall and FUGAKU), and HPCI system (Project ID: hp210107, hp230090). A.N. was funded by a Riken Special Postdoctoral Researcher Fellowship, Grants-in-Aid for JSPS Fellow (21J40162), and JST PRESTO (JPMJPR22A9). A.N, J.J. and Y.S. were funded by JSPS KAKENHI (17K18365, 21H00412, 22H05438 and 24H01162 (to A.N.), 21H05282 (to J.J.), 21H05249 (to Y.S.)). A.N. and K.R.M. were supported by a BBSRC grant to H.B. and D.N.W. (BB/J009784/1). A.J.S. was funded by the Bristol Chemical Synthesis Centre for Doctoral Training funded by the EPSRC (EP/G036764/1). A.N., A.J.S., A.R.T., and D.N.W. were funded by ERC Grants to D.N.W. (340764 and 787173). M.I.W. was funded by the BBSRC (BB/R001790/1). D.N.W. held a Royal Society Wolfson Research Merit Award (WM140008).

## Notes

### Competing Interest Statement

The authors have declared no competing interest.

