## Supporting Information for "De novo design of α-helical peptide channels with designer stoichiometry"

### Contents

1. Supplementary Methods ........ S3

2. Supplementary Tables ........ S5

3. Supplementary Figures ........ S8

**Supplementary Methods**

**Materials**

The following materials were used for this study: Fmoc-L-amino acids, dimethylformamide (DMF) and activators (AGTC Bioproducts), Rink amide ChemMatrixTM resin (PCAS Biomatrix Inc), Boc-Ser(Fmoc-Ala)-OH and Boc-Thr(Fmoc-Ala)-OH (Novabiochem, Merck Millipore, Germany), 1,2-diphytanoyl-*sn*-glycero-3-phosphocholine (DPhPC stored as a chloroform stock at 50 mg/mL at -20 C) (Avanti Polar Lipids), $Ca^{2+}$-sensitive dye Fluo-8H (AAT Bioquest, California, USA), all other solvents (Fisher Scientific). All other substances were purchased from Sigma-Aldrich unless stated otherwise, and all aqueous solutions were made using 18.2 MΩ.cm Milli-Q water.

**Peptide synthesis**

All peptides were synthesized by the Fmoc solid-phase peptide synthesis with a CEM Liberty Blue automated synthesizer with inline UV monitoring. N,N'-diisopropylcarbodiimide/1-hydroxybenzotriazole (1/1 molar ratio) was used as an activator. All peptides were synthesized using Rink amide resin, yielding a C-terminal amide, and were N-terminally acetylated using N,N-diisopropylethylamine (4.5 eq.) and acetic anhydride (3 eq.) in N,N-dimethylformamide (DMF) for 30 min. Boc-Ser(Fmoc-Ala)-OH and Boc-Thr(Fmoc-Ala)-OH were used at the A10-S/T20. The isoacyl dipeptide block was manually coupled with the activator mixture in dichloromethane/DMF (3/1, v/v) for one hour. The completion of the coupling reaction was monitored by a 2,4,6-trinitrobenzenesulfonic acid test, and the coupling was repeated up to three times as required. The rest of the peptide after the dipeptide block was synthesized by using the synthesizer without microwave heating. Cleavage from the solid support was carried out with a mixture of trifluoroacetic acid (TFA)/triisopropylsilane/water (90/5/5, v/v/v). An additional 5% ethanedithiol was added to the cleavage mixture for all Cys-containing peptides. The cleavage solution was reduced in volume to ~1 mL by using a flow of nitrogen. Diethyl ether (45 mL) was added to obtain a precipitate that was recovered by centrifugation. The precipitate was dissolved in acetonitrile/water (1/1) before freeze-drying, to obtain the crude peptide. Peptides were purified by reversed-phase HPLC, with a gradient of 30 to 80% acetonitrile in water (each containing 0.1% TFA) over 45 min on a Vydac ® TP C18 column (10 μm particle, 22 × 250 mm), followed by the second purification with a gradient of 30 to 60% acetonitrile in water over 35 min on a Phenomenex® Luna C8 column (5 μm particle, 10 mm × 250 mm). Fractions containing pure peptide were identified by analytical HPLC and MALDI-TOF mass spectrometry. The purified O-acyl peptides were subsequently rearranged to N-acyl peptides, where 50 μM peptide in trifluoroethanol (TFE)/water (1/1, v/v) was treated with 100 mM ammonium bicarbonate. The reaction was monitored using analytical HPLC and when completed (times ranged from 1h – overnight) the peptide solution was freeze-dried and redissolved in hexafluoroisopropanol/water. Freeze-drying and dissolving were repeated four times to ensure the removal of the ammonium bicarbonate.

Analytical HPLC was performed with a Jasco 2000 series HPLC system with a Phenomenex Kinetex C18 column (5 µm particle, 4.6 mm × 100 mm) for fraction identification and Phenomenex Aeris Widepore C4 column (3.6µm particle, 4.6 mm × 100 mm) for reaction monitoring, monitored at 220 nm. The gradient was 20 to 80% acetonitrile/water (each containing 0.1% TFA) over 16 min for the C18 column and 40 to 95% over 20 min for the C4 column. MALDI-TOF mass spectra were obtained on a Bruker UltraFlex MALDI-TOF mass spectrometer in positive-ion reflector mode. Peptide solutions were spotted on a ground steel target plate with dihydroxybenzoic acid as the matrix. Calibration was conducted using the 'nearest neighbor' method, with Bruker Peptide Calibration Standard II as the reference masses.

#### Supplementary Tables

**Table S1.** bZIP scoring for the heptad sequences selected for synthesis. Interface pairs ***gade*1** and ***gade*2** correspond to positions ***cdga*** and ***deab***, respectively. Fitness scores were calculated by subtracting the higher Self-association score from Raw scores.

| **Sequence** *gabcdef* | Fitness score | Raw score | gade1 Self-association score | gade2 Self-association score |
|---|---|---|---|---|
| **ASVINAx** | 49.652 | 30.987 | -24.165 | -18.665 |
| **ATVINAx** | 33.044 | 26.99 | -6.054 | -16.675 |
| **ASVANAx** | 45.989 | 27.324 | -41.376 | -18.665 |

**Table S2.** Complete sequences of synthesized peptides.

| Name | N-term | Solubility tag | Coiled-coil sequence | C-term |
|---|---|---|---|---|
| | | | **cdefgabcdefgabcdefgabcdefgab** | |
| CCTM-$S_aV_bI_cN_d$ | Ac- | KKKKGSG | INAWASVINALASVINALASVINAWASV | G-$NH_2$ |
| CCTM-$T_aV_bI_cN_d$ | Ac- | KKKKGSG | INAWATVINALATVINALATVINAWATV | G-$NH_2$ |
| CCTM-$S_aV_bA_cN_d$ | Ac- | KKKKGSG | ANAWASVANALASVANALASVANAWASV | G-$NH_2$ |
| CCTM-$S_aV_bI_cN_d$-S20C | Ac- | KKKKGSG | INAWASVINALASVINALACVINAWASV | G-$NH_2$ |
| CCTM-$S_aV_bI_cN_d$ [KLLW] | Ac- | KKKKGSG | INAKASVINALASVINALASVINAWASV | G-$NH_2$ |

**Table S3.** Crick parameters and BUDE scores of the coiled-coil barrel models optimized in ISAMBARD.

| | Model | | | | | | Experiment |
|---|---|---|---|---|---|---|---|
| | Oligomeric state | BUDE score /chain | Number of residues | Radius (Å) | Pitch (Å) | Interface angle (°) | Oligomer state |
| CCTM-$S_aV_bI_cN_d$ | 4 | -46.8 | 30 | 6.37 | 217 | 75.8 | 5 |
| | 5 | -65.21 | 30 | 8.15 | 178 | 116.49 | |
| | 6 | -85.21 | 30 | 8.68 | 270 | 115.80 | |
| | 7 | -73.80 | 30 | 11.28 | 201 | 129.91 | |
| | 8 | -112.27 | 30 | 11.77 | 201 | 119.24 | |
| CCTM-$S_aV_bA_cN_d$ | 5 | -94.31 | 30 | 7.79 | 196 | 115.85 | 7 |
| | 6 | -125.16 | 30 | 8.77 | 217 | 116.60 | |
| | 7 | -138.83 | 30 | 9.77 | 500 | 115.41 | |
| | 8 | -153.97 | 30 | 11.95 | 201 | 116.26 | |
| CCTM-$T_aV_bI_cN_d$ | 5 | -100.55 | 30 | 7.70 | 291 | 115.00 | 7 |
| | 6 | -130.04 | 30 | 8.61 | 439 | 115.00 | |
| | 7 | -153.52 | 30 | 9.77 | 498 | 115.41 | |
| | 8 | -154.42 | 30 | 11.43 | 495 | 115.46 | |

**Table S4.** Summary of MD simulation systems

| peptide | CCTM-$S_aV_bI_cN_d$ | CCTM-$S_aV_bI_cN_d$ | CCTM-$S_aV_bI_cN_d$ | CCTM-$S_aV_bI_cN_d$ | CCTM-$S_aV_bI_cN_d$ |
|---|---|---|---|---|---|
| # of peptides | 4 | 5 | 6 | 7 | 8 |
| # of lipids | 160 | 161 | 160 | 161 | 161 |
| # of atoms | 52741 | 59991 | 54243 | 63794 | 54930 |
| Box size (x, y, z) (Å) | 83, 83, 82 | 85, 85, 80 | 85, 85, 80 | 86, 86, 83 | 87, 87, 78 |

| peptide | CCTM-$S_aV_bI_cN_d$ | CCTM-$S_aV_bI_cN_d$ | CCTM-$S_aV_bI_cN_d$ | CCTM-$S_aV_bI_cN_d$-S20Cace | CCTM-$T_aV_bI_cN_d$ | CCTM-$S_aV_bA_cN_d$ |
|---|---|---|---|---|---|---|
| # of peptides | 10 | 15 | 20 | 6 | 7 | 7 |
| # of lipids | 399 | 399 | 399 | 161 | 160 | 160 |
| # of atoms | 161322 | 174261 | 188298 | 66770 | 60006 | 59371 |
| Box size (x, y, z) (Å) | 133, 133, 96 | 138, 138, 97 | 144, 144, 97 | 87, 87, 87 | 86, 86, 87 | 85, 85, 87 |

**Table S5.** Summary of equilibration steps before production runs of MD simulations

| Step | 0 | 1 | 2 | 3 | 4 | 5 | 6 |
|---|---|---|---|---|---|---|---|
| # of steps | 20000 | 25000 | 25000 | 25000 | 50000 | 50000 | 50000 |
| Time step (fs) | | 1.0 | 1.0 | 1.0 | 2.0 | 2.0 | 2.0 |
| Ensemble | Energy minimization | NVT | NVT | NPT | NPT | NPT | NPT |
| **Positional restraint (kcal/mol)** | | | | | | | |
| Peptide backbone | 10.0 | 10.0 | 5.0 | 2.5 | 1.0 | 0.5 | 0.1 |
| Peptide sidechain | 5.0 | 5.0 | 2.5 | 1.0 | 0.5 | 0.1 | 0 |
| Lipid P (Z direction only) | 2.5 | 2.5 | 2.5 | 1.0 | 0.5 | 0.1 | 0 |

#### Supplementary Figures

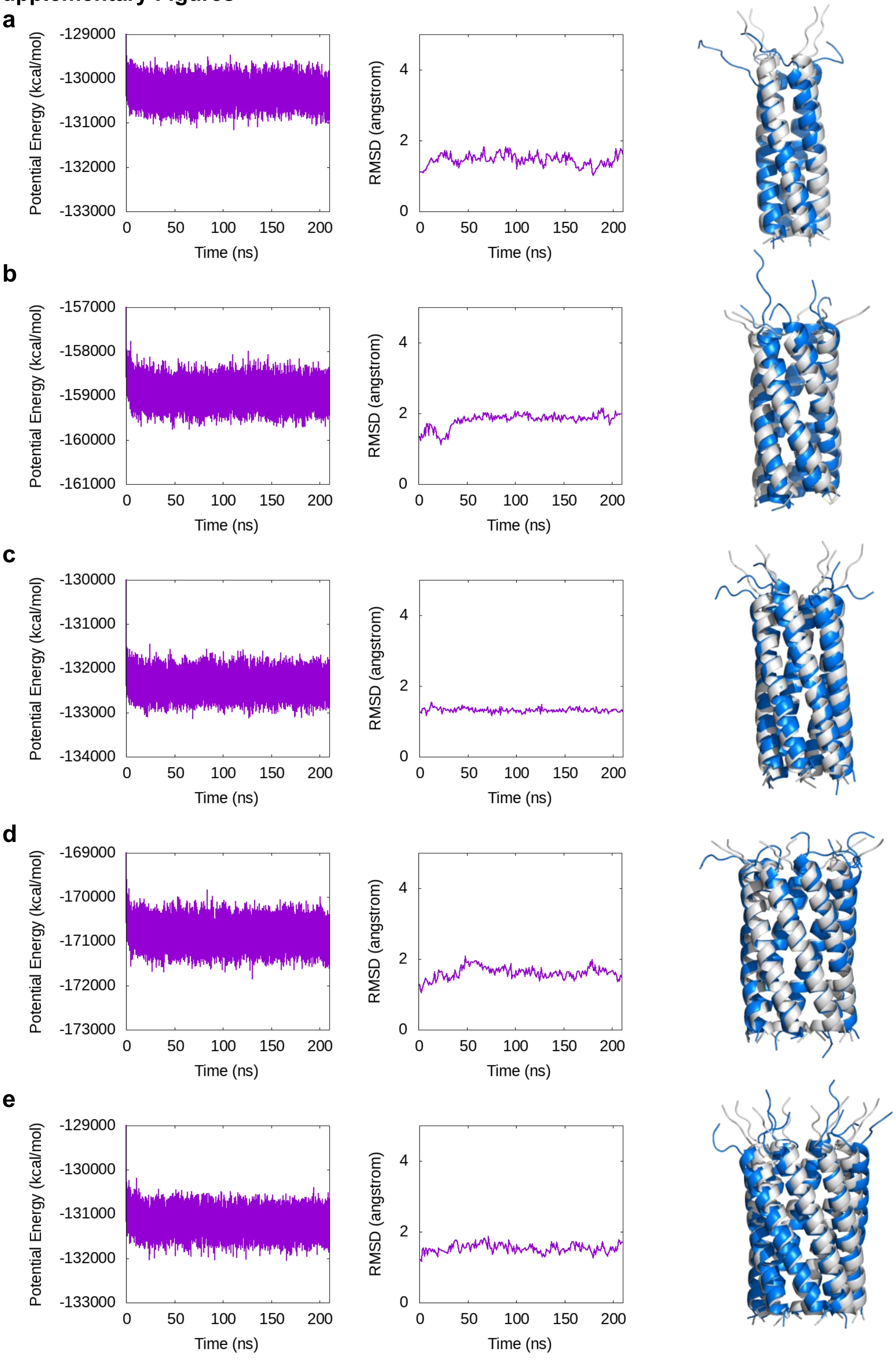

**Figure S1.** All-atom MD simulations of CCTM-$S_aV_bI_cN_d$ barrel models in DPhPC lipid bilayer. Time course of potential energy (left), RMSD of backbone α carbons of residues 8–35 with respect to the initial model and (center), and overlayed structures (right) of initial models (gray) and snapshots after 200 ns MD simulations (blue). (a) Tetramer, (b) pentamer, (c) hexamer, (d) heptamer, and (e) octamer.

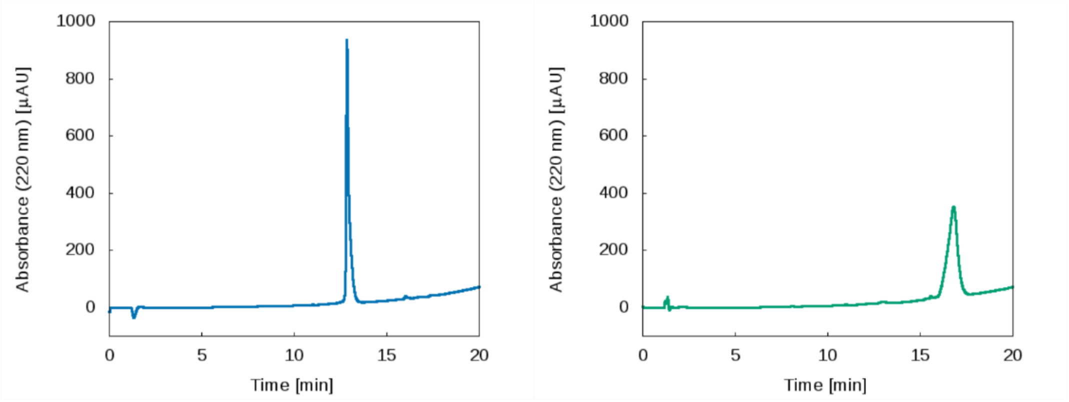


**Figure S2.** HPLC traces of CCTM-$S_aV_bI_cN_d$-O (left) and CCTM-$S_aV_bI_cN_d$-N (right) from a linear gradient of 40 to 95% acetonitrile/water (0.1 % TFA)

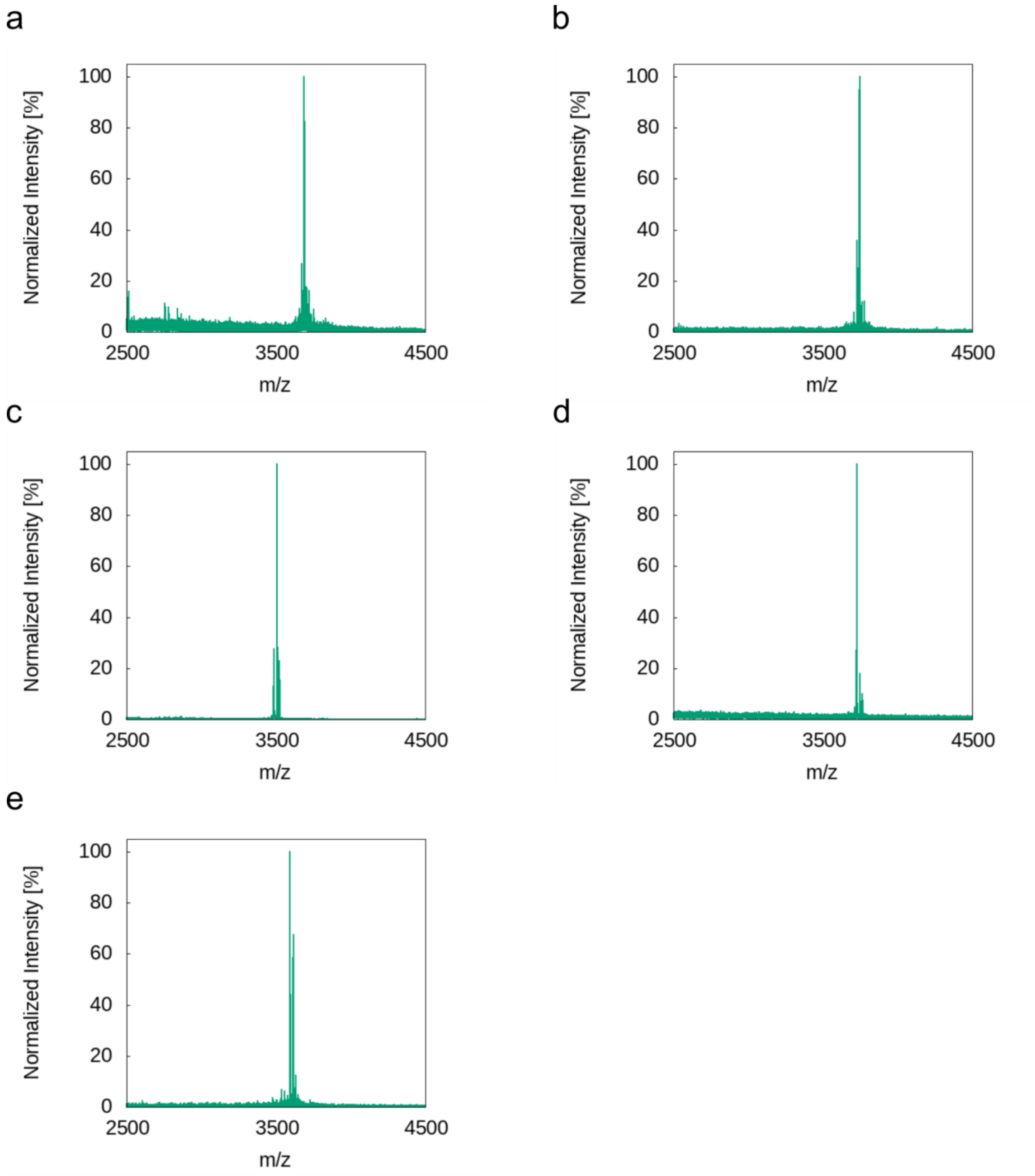


**Figure S3.** MALDI-TOF MS spectra. (a) CCTM-$S_aV_bI_cN_d$, Calculated mass = 3650.043Da, Observed mass = 3650.808 Da, (b) CCTM-$T_aV_bI_cN_d$, Calculated mass = 3706.106 Da, Observed mass = 3705.929 Da, (c) CCTM-$S_aV_bA_cN_d$, Calculated mass = 3481.855 Da, Observed mass = 3481.954 Da, (d) CCTM-$S_aV_bI_cN_d$-S20C-ace, Calculated mass = 3724.060 Da, Observed mass = 3724.935 Da, (e) CCTM-$S_aV_bI_cN_d$-KLLW, Calculated mass = 3592.059 Da, Observed mass = 3591.921 Da

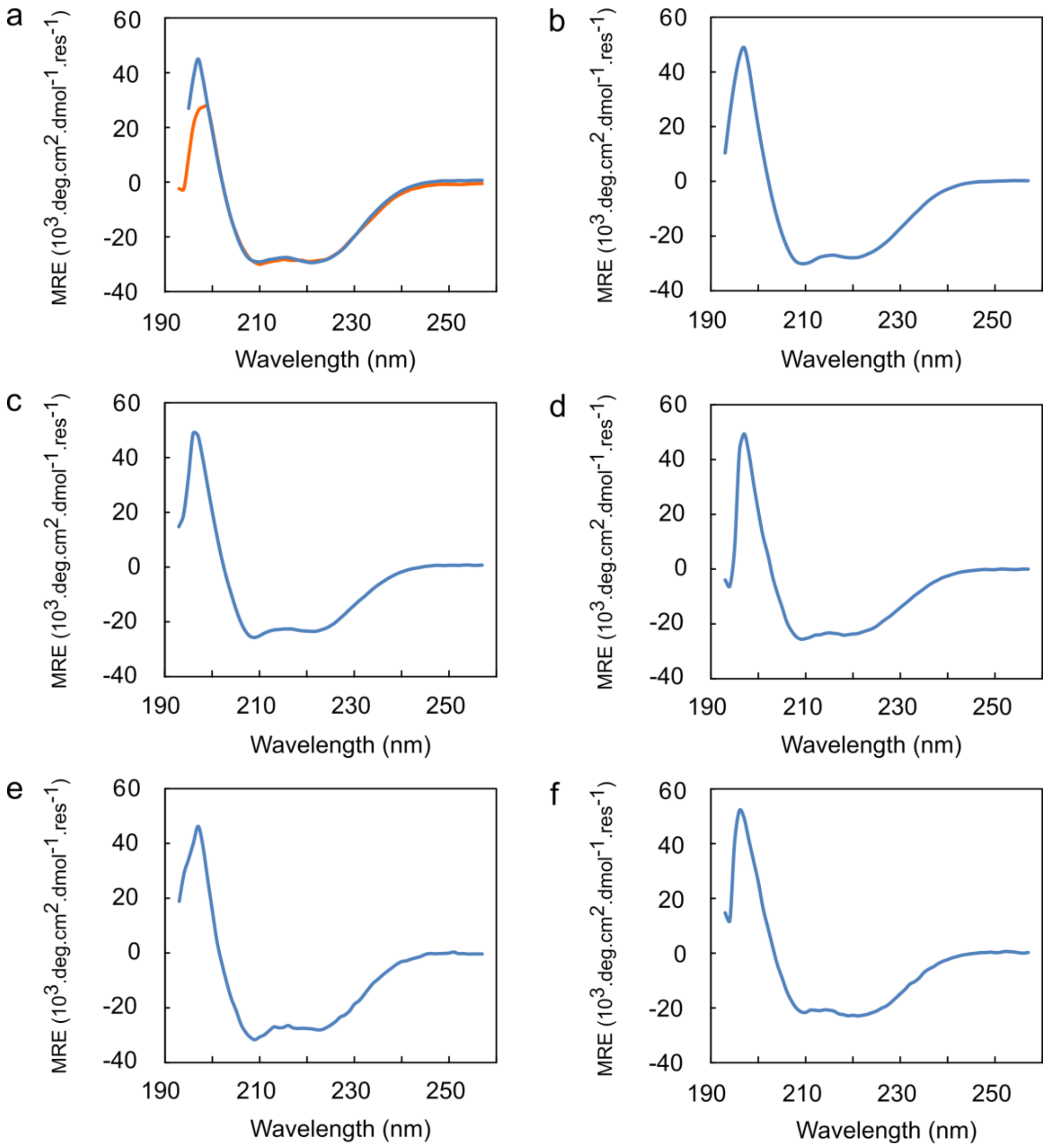


**Figure S4.** (a) CD spectra measured at 20°C with 20–25 μM peptide in PBS (pH7.4) with 0.35% C8E5 (blue) and with 1.5% OG (orange) for CCTM-$S_aV_bI_cN_d$, (b-f) CD spectra measured at 20°C with 20–25 μM peptide in PBS (pH7.4) with 0.35% C8E5 for CCTM-$T_aV_bI_cN_d$ (b), CCTM-$S_aV_bA_cN_d$ (c), CCTM-$S_aV_bI_cN_d$-S20C (d), CCTM-$S_aV_bI_cN_d$-S20C-ace (e), and CCTM-$S_aV_bI_cN_d$-KLLW (f).

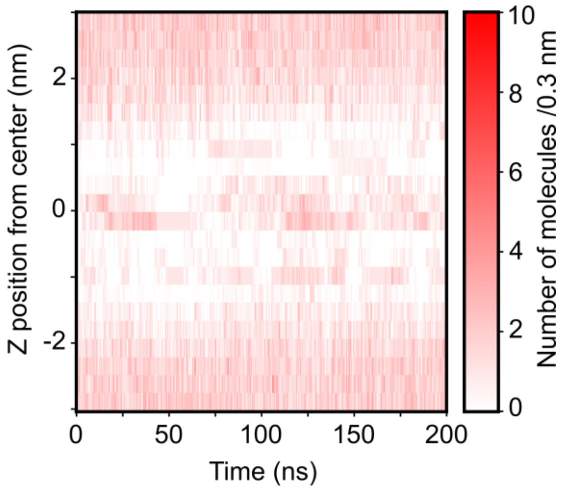


**Figure S5.** Water permeation through the pentameric CCTM-$S_aV_bI_cN_d$ channel. Z positions of oxygen atoms of 57 water molecules that accessed the central region of the lumen (-0.5 to +0.5 nm) during the MD simulation. Molecules within every 0.3 nm were counted and plotted every 10 ps.

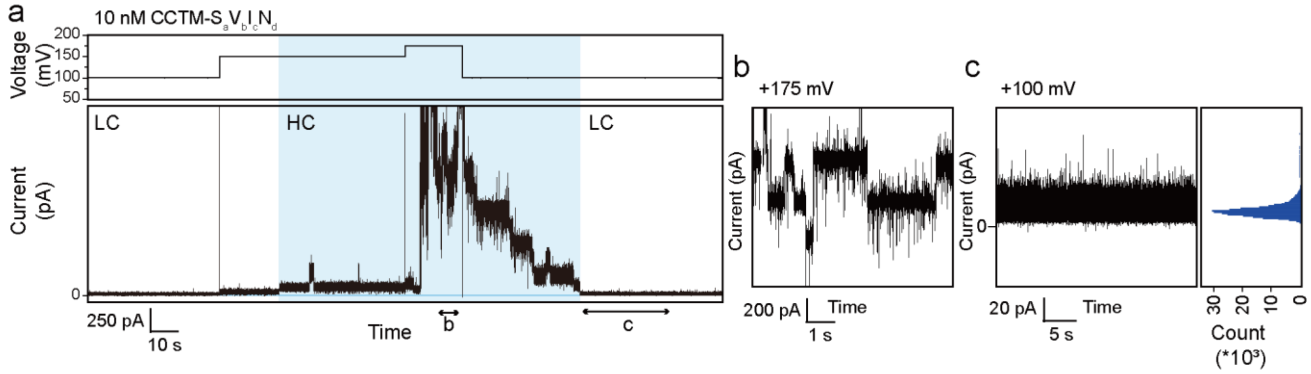


**Figure S6.** Voltage-dependent transition between LC and HC states of CCTM-$S_aV_bI_cN_d$. (a) Representative current recording (lower box) of CCTM-$S_aV_bI_cN_d$ (10 nM) with transitions between the LC and HC state dependent on applied voltages (upper box). The HC state is highlighted with light blue. (b) Expansion of the range "b" in the recording (a) showing the HC state. (c) Expansion of the range "c" in the recording (a) showing the LC state.

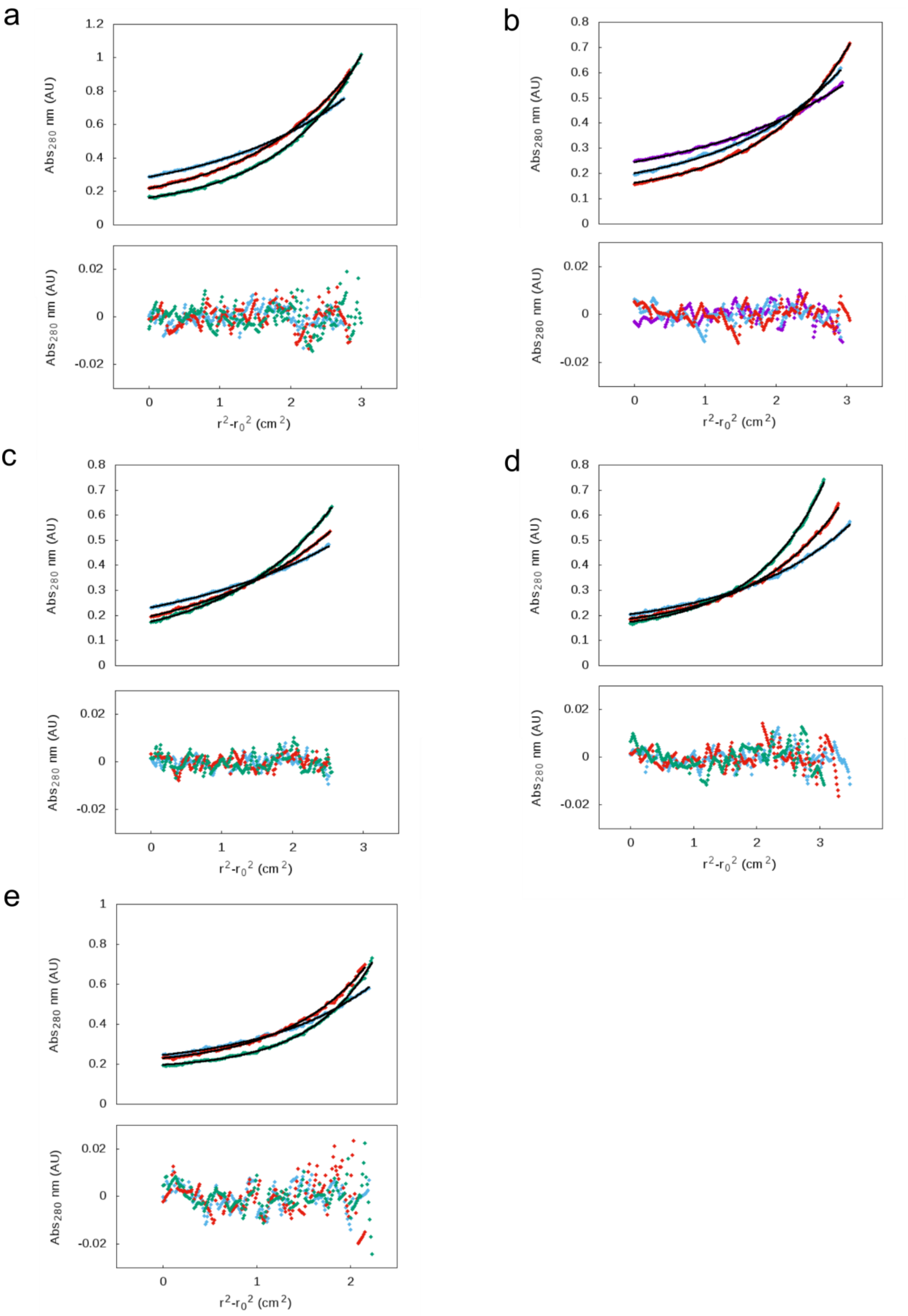

a
b
c
d
e
Abs280 nm (AU)
r2-r02 (cm2)

**Figure S7.** Analytical ultracentrifugation sedimentation equilibrium data. Sedimentation equilibrium data (top, dots), fitted single-component equilibrium model curves (top, lines) and residuals (bottom). Three rotor speeds were selected out of 18k (purple), 21k (blue), 24k (red), and 27k (green) rpm. Conditions: 20–25 $\mu$M peptide, 20°C, PBS, 0.35% C8E5.
(a) CCTM-$T_aV_bI_cN_d$, the fits return a mass of 25969 Da (7.0 × monomer mass, 95% confidence limits 25529 – 26420 Da). (b) CCTM-$S_aV_bA_cN_d$, the fits return a mass of 24487 Da (7.0 × monomer mass, 95% confidence limits 24214 – 24761 Da). (c) CCTM-$S_aV_bI_cN_d$ S20C, the fits return a mass of 17093 Da (4.7 × monomer mass, 95% confidence limits 16772 – 17373 Da). (d) CCTM-$S_aV_bI_cN_d$-S20C-ace, the fits return a mass of 22838 Da (6.1 × monomer mass, 95% confidence limits 22350 – 23317 Da). (e) CCTM-$S_aV_bI_cN_d$ [KLLW], the fits return a mass of 38380 kDa = ~11 x monomer mass, although its stoichiometry was not conclusive due to the poor fit to the single-component model.

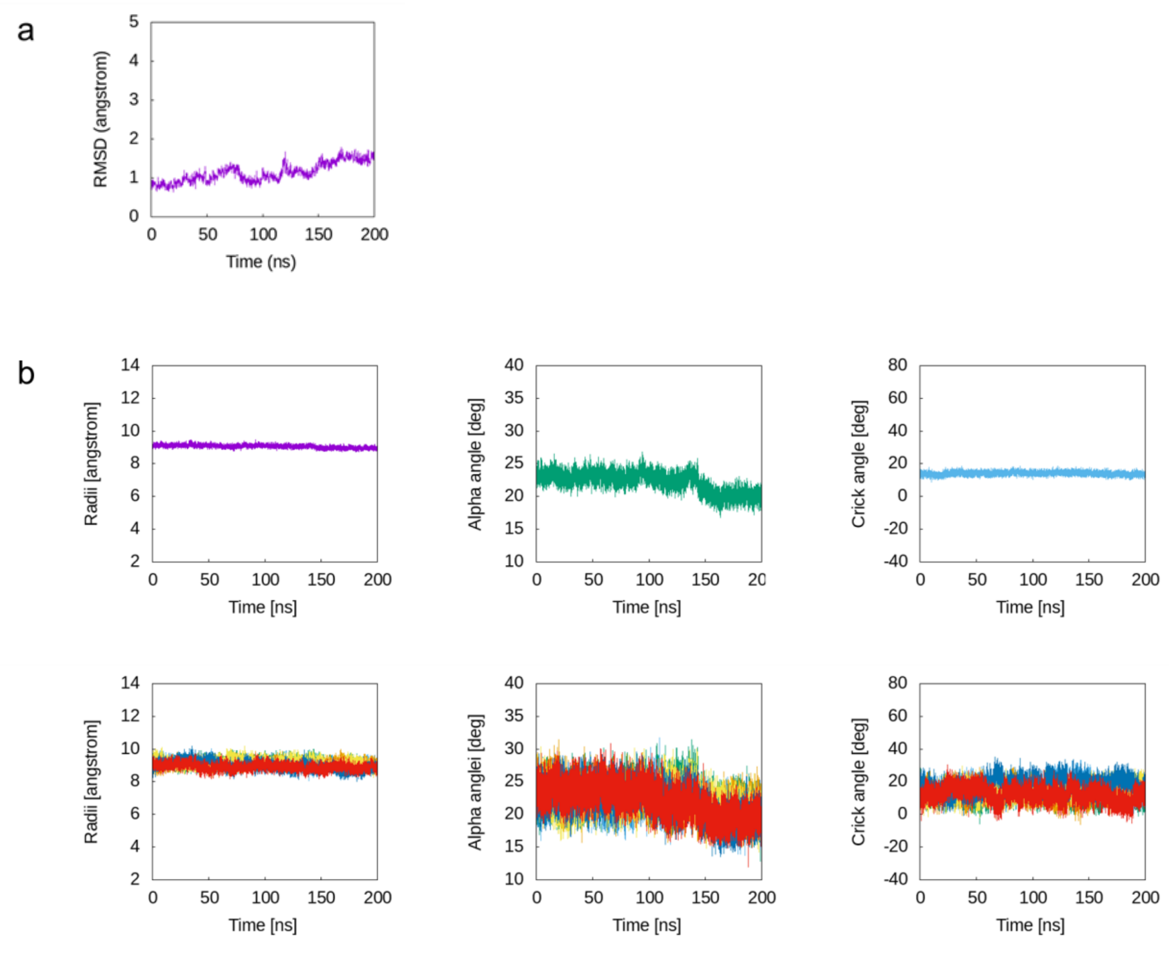


**Figure S8.** Time courses of RMSD and coiled-coil Crick parameters of CCTM-$S_aV_bI_cN_d$-S20C-ace hexamer in 200-ns MD simulations. (a) Time course of RMSD of C$\alpha$ excluding the Lys tail region (b) Time courses of the average of all helices (upper) and each chain (lower) of Helix radius (left column), Alpha angle (middle column), and Crick angle (right column)

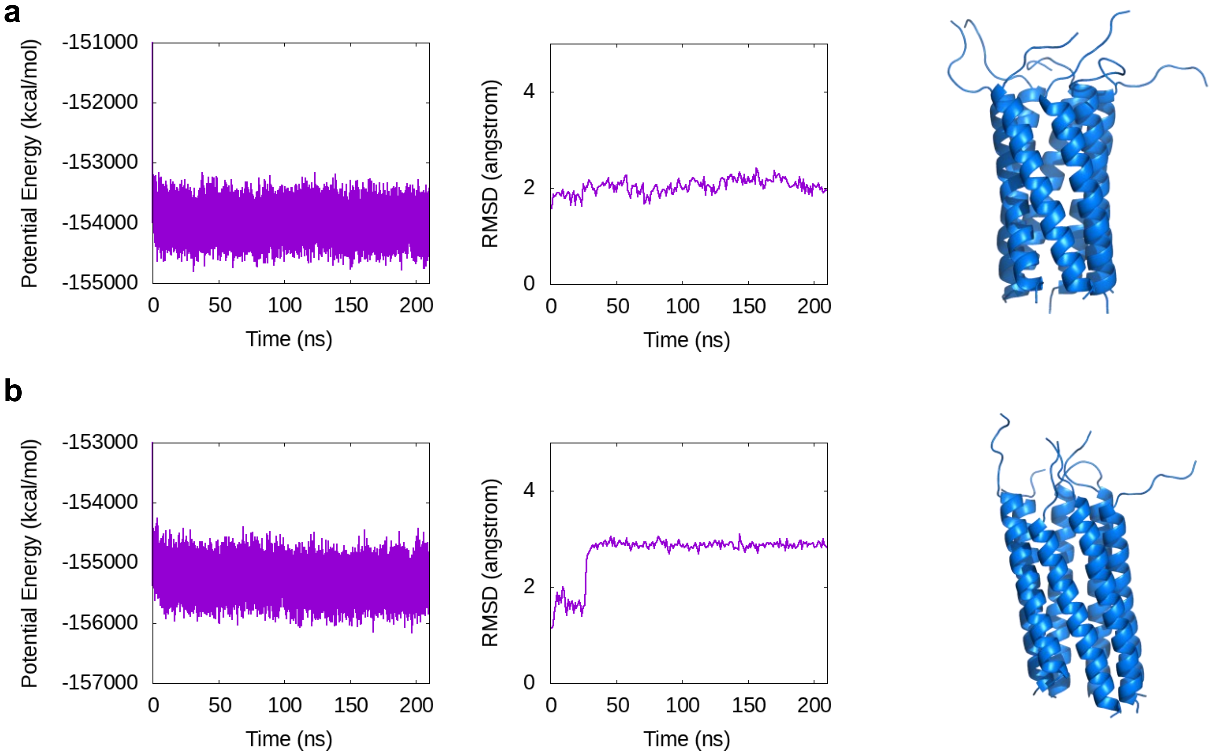


**Figure S9.** All-atom MD simulations of heptameric CCTM-$S_aV_bA_cN_d$ (a) and CCTM-$T_aV_bI_cN_d$ (b) models in DPhPC lipid bilayer. Time course of potential energy (left), RMSD of backbone α carbons of residues 8–35 with respect to the initial model and (center), and snapshots after 200 ns MD simulations (right).

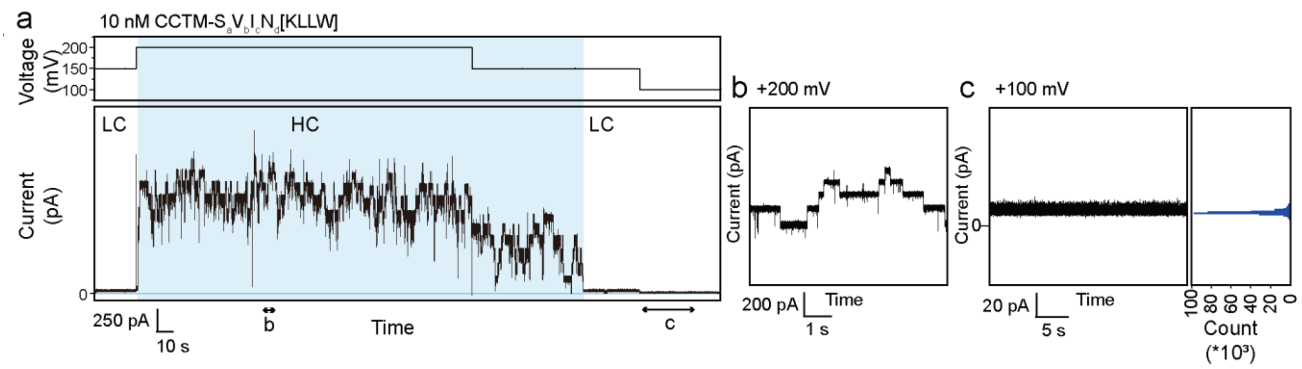


**Figure S10.** Voltage-dependent transition between LC and HC states of CCTM-$S_aV_bI_cN_d$. (a) Representative current recording (lower box) of CCTM-$S_aV_bI_cN_d$ (10 nM) with transitions between the LC and HC state dependent on applied voltages (upper box). The HC state is highlighted with light blue. (b) Expansion of the range "b" in the recording (a) showing the HC state. (c) Expansion of the range "c" in the recording (a) showing the LC state.

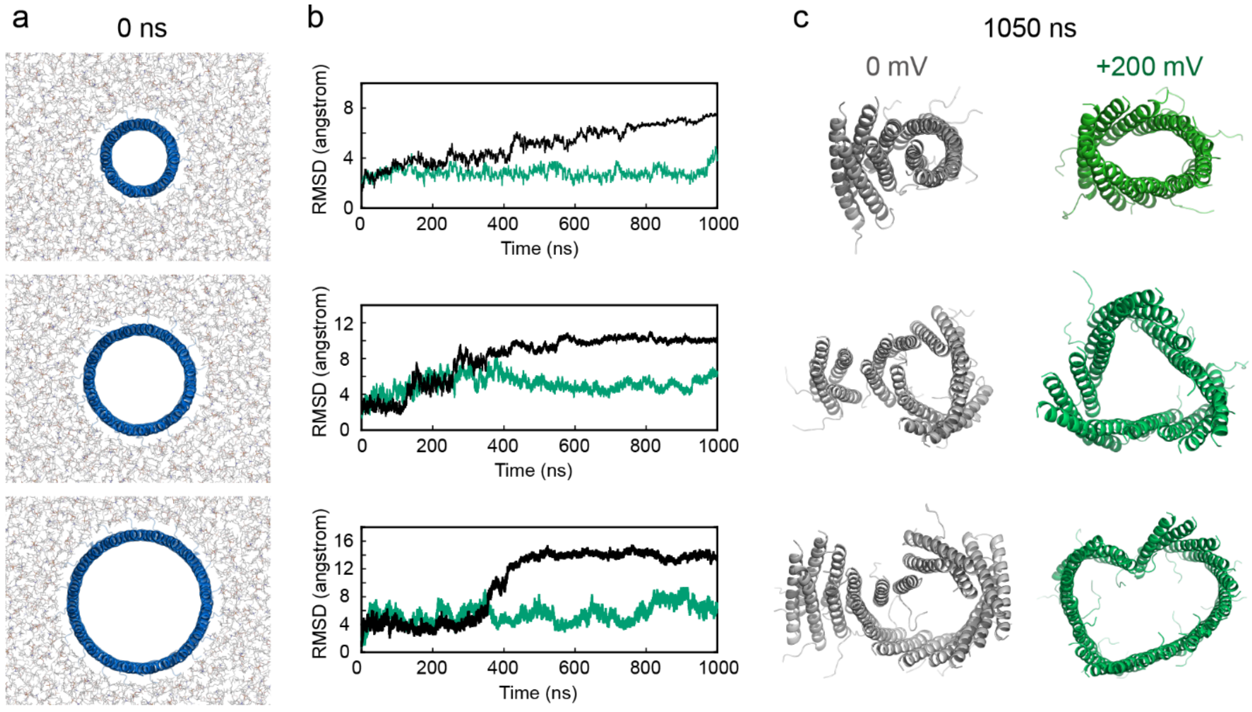


**Figure S11**. All-atom molecular dynamics simulations of CCTM-$S_aV_bI_cN_d$ barrels. (a) Initial configurations of 10- (top), 15- (middle), and 20-mer (bottom) barrels embedded in a 1,2-diphytanoyl-sn-glycero-3-phosphocholine bilayer viewed from the C-terminal side of the peptides. (b) Time course of RMSD of Cα of 10- (top), 15- (middle), and 20-mer (bottom) barrels. Black and green lines correspond to simulations without an external electric field and with +200 mV potential, respectively. (c) Final snapshots of 10- (top), 15- (middle), and 20-mer (bottom) barrels without an external electric field (left, black cartoon) and with +200 mV potential (right, green cartoon).

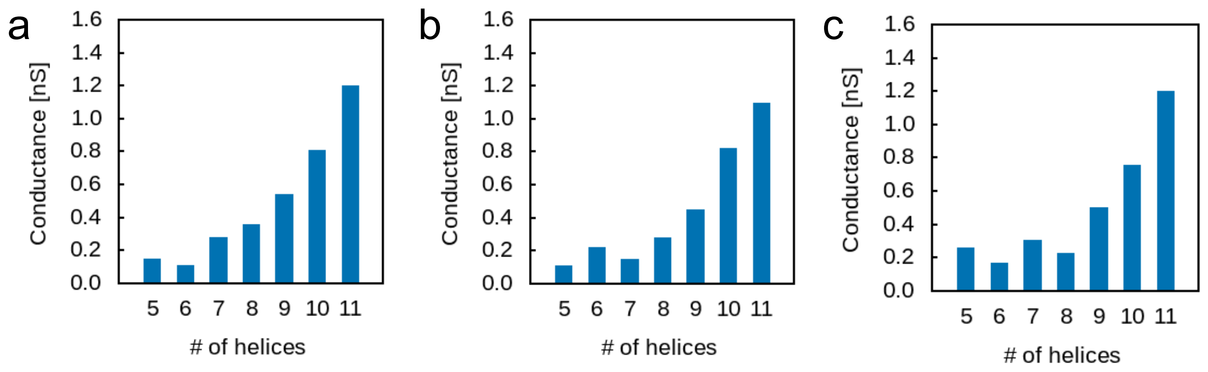


**Figure S12**. Predicted conductance of optimized coiled-coil barrel models. Estimated conductance by HOLE program of (a) CCTM-S$_a$V$_b$I$_c$N$_d$, (b) CCTM-T$_a$V$_b$I$_c$N$_d$, and (c) CCTM-S$_a$V$_b$A$_c$N$_d$ with 1M KCl.

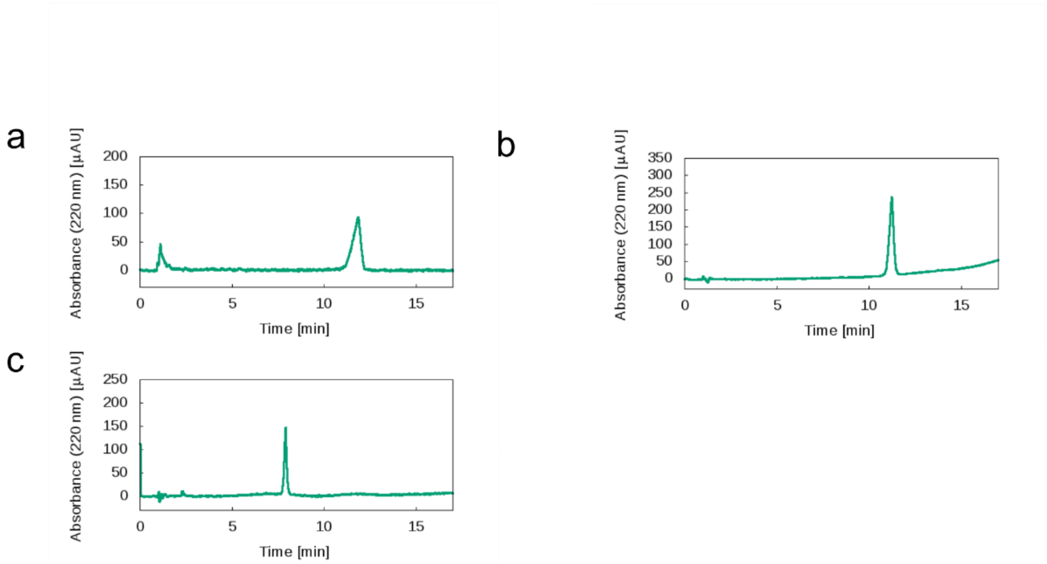


**Figure S13.** HPLC traces of CCTM-T$_a$V$_b$I$_c$N$_d$ (a), CCTM-S$_a$V$_b$A$_c$N$_d$ (b), and CCTM-S$_a$V$_b$I$_c$N$_d$-KLLW (c) from a linear gradient of 20 to 80% acetonitrile/water (each containing 0.1% TFA), monitored at 220 nm.

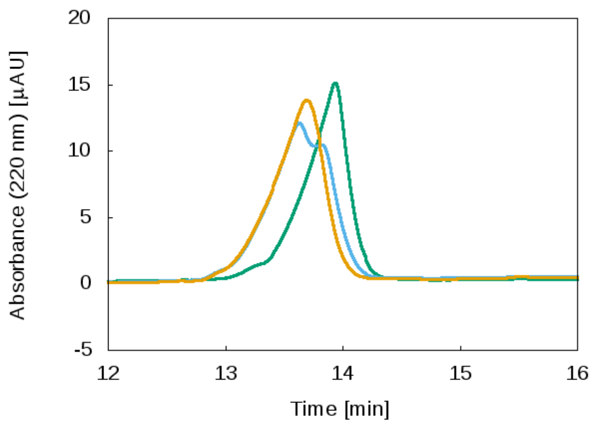


**Figure S14.** HPLC traces of CCTM-$S_aV_bI_cN_d$-S20C (green), alkylation reaction solution after 1h (blue), and CCTM-$S_aV_bI_cN_d$-S20C-ace (orange) from a linear gradient of 40 to 95 % acetonitrile/water (0.1%TFA) over 30 min.

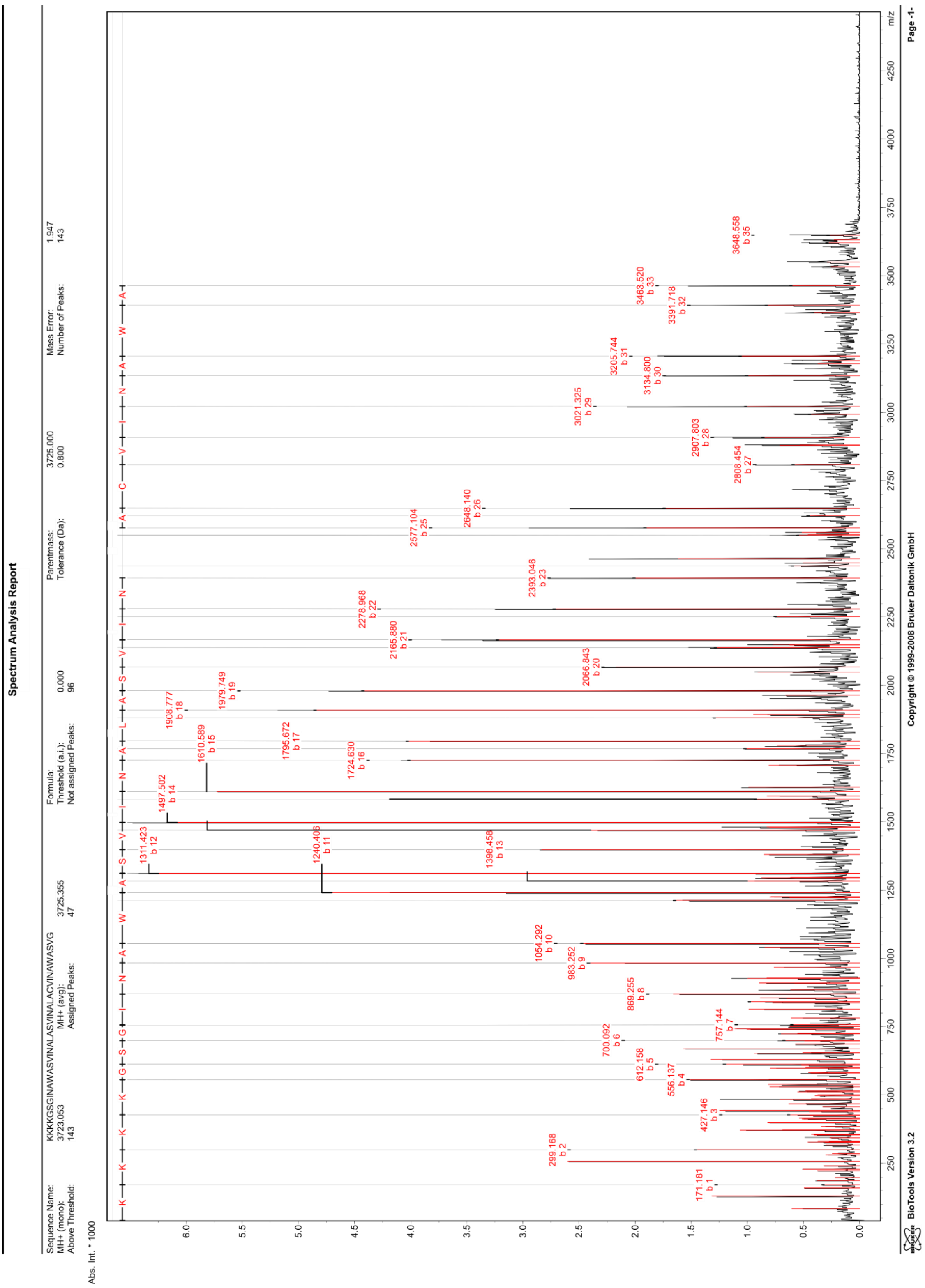


**Figure S15.** MALDI-TOF MS/MS spectrum of CCTM-$S_aV_bI_cN_d$-S20C-ace with peak assignments. The successful alkylation of the cysteine residue was confirmed.

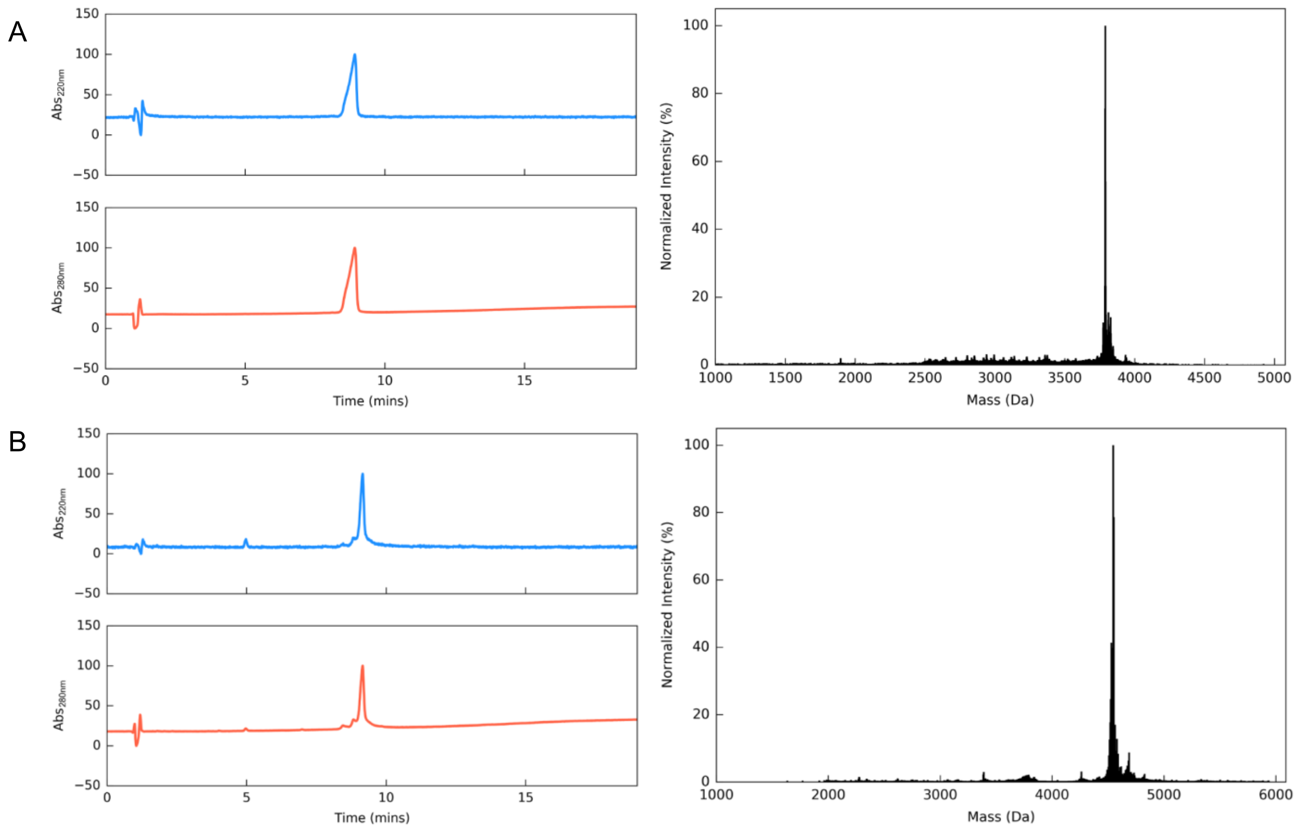


**Figure S16.** (A) Cys-Gly-CCTM-$S_aV_bI_cN_d$ [KLLW]: HPLC traces (left) from a linear gradient of 40 to 100% MeCN (0.1% TFA) in H2O (0.1% TFA), monitored at 220 nm (blue) and 280 nm (red). MALDI-TOF mass spectrum (right). Calculated monoisotopic peptide mass = 3808.1 Da, observed mass = 3809 Da. (B) Cy5-labeled CCTM-$S_aV_bI_cN_d$ [KLLW]: HPLC traces (left) from a linear gradient of 40 to 100% MeCN (0.1% TFA) in $H_2O$ (0.1% TFA), monitored at 220 nm (blue) and 280 nm (red). Negative mode MALDI-TOF mass spectrum (right). Calculated monoisotopic peptide mass = 4572.0, observed mass = 4571 Da.
